# Group 2 innate lymphoid cells promote inhibitory synapse development and social behavior

**DOI:** 10.1101/2023.03.16.532850

**Authors:** Jerika J. Barron, Nicholas M. Mroz, Sunrae E. Taloma, Madelene W. Dahlgren, Jorge Ortiz-Carpena, Leah C. Dorman, Ilia D. Vainchtein, Caroline C. Escoubas, Ari B. Molofsky, Anna V. Molofsky

## Abstract

The innate immune system plays essential roles in brain synaptic development, and immune dysregulation is implicated in neurodevelopmental diseases. Here we show that a subset of innate lymphocytes (group 2 innate lymphoid cells, ILC2s) is required for cortical inhibitory synapse maturation and adult social behavior. ILC2s expanded in the developing meninges and produced a surge of their canonical cytokine Interleukin-13 (IL-13) between postnatal days 5-15. Loss of ILC2s decreased cortical inhibitory synapse numbers in the postnatal period where as ILC2 transplant was sufficient to increase inhibitory synapse numbers. Deletion of the IL-4/IL-13 receptor (*Il4ra*) from inhibitory neurons phenocopied the reduction inhibitory synapses. Both ILC2 deficient and neuronal *Il4ra* deficient animals had similar and selective impairments in adult social behavior. These data define a type 2 immune circuit in early life that shapes adult brain function.

**One sentence summary:** Type 2 innate lymphoid cells and Interleukin-13 promote inhibitory synapse development.

## INTRODUCTION

The innate immune system monitors and promotes healthy organ development, including in the brain. Conversely, immune dysfunction has both epidemiologic and genetic links to neurodevelopmental and psychiatric disorders (Bohlen et al., 2019; Bennett and Molofsky, 2019; Knuesel et al., 2014). An increase in the balance of excitatory to inhibitory synapses during development is a conserved feature of neurodevelopmental disorders including autism spectrum disorder and epilepsy (Rubenstein, 2010; Rubenstein and Merzenich, 2003). However, the mechanisms of this synaptic imbalance are likely multifactorial. One potential immune link to behavior and cognition includes lymphocytes and lymphocyte-derived cytokines (Derecki et al., 2010; Filiano et al., 2016; Kipnis, 2016; Brombacher et al., 2017; Ribeiro et al., 2019; Alves de Lima et al., 2020; Herz et al., 2021). Defining the cellular circuits through which lymphocytes shape the developing brain is thus essential to understanding their impacts in neurodevelopmental diseases.

Homeostatic immune responses occur locally within tissues and are predominantly innate responses in early life. Innate immune cells including macrophages and innate lymphoid cells (ILCs) are deposited during embryogenesis and early postnatal life. ILCs locally mature their effector functions concomitant with organ maturation (Gasteiger et al., 2015). Innate immune responses are particularly critical in this early life window, where they can act prior to the full maturation and expansion of adaptive B and T cells. ILCs are the most recently discovered member of the innate immune arsenal (Kotas and Locksley, 2018). Their subtypes closely mirror the types of T cells that expand later in life (*i.e*. type 1, type 2, type 3/17). Unlike many T cells, ILCs produce cytokines within hours and in response to local tissue cues, rather than in response to specific antigens. Group 2 innate lymphoid cells (ILC2s) represent one of these subsets and are dominant producers of the type 2 cytokine Interleukin-13 (IL-13), through which they drive tissue remodeling. This is critical to limiting helminths and driving allergic pathology, but also for tissue development and remodeling (Kobayashi et al., 2019; Saluzzo et al., 2017). In the brain, type 2 immune signals including Interleukin-33 drive excitatory synaptic remodeling in brain development and plasticity (Vainchtein et al., 2018; Nguyen et al., 2020; Han et al., 2023).

Here we show that ILC2s and their canonical cytokine IL-13 promote inhibitory synapse development in early life and are required for adult social behavior. We found that ILC2s expand perinatally in the mouse brain meninges and produce a burst of IL-13 between postnatal days 5-15, during a key window of neuronal synapse maturation. ILC2-deficient mice had a reduction in cortical inhibitory synapses at postnatal day 15 (P15), and interneuron-specific deletion of the IL-13/IL-4 receptor subunit *Il4ra* phenocopied these synaptic deficits. Conversely, intracranial injection of ILC2s or of IL-13 was sufficient to increase inhibitory synapse numbers. Finally, both ILC2 deficiency or interneuron-specific *Il4ra* deficiency led to impaired social interaction in adulthood. These data support a model in which ILC2s signal to interneurons via IL-13 to promote inhibitory synapse maturation. More broadly, they suggest that innate immune signaling can dynamically shape neuronal connectivity during key periods of brain development.

## RESULTS

### Meningeal type 2 innate lymphocytes expand and produce IL-13 in early life

The CNS immune environment includes cells within the brain parenchyma (primarily microglia) as well as in the brain boundaries, which include the meninges and perivascular spaces. These boundary regions contain a full complement of immune cell types, including macrophages, dendritic cells, and innate and adaptive lymphocytes that mirror tissue immune niches in other organs (Rua and McGavern, 2018; Alves De Lima et al., 2020). To define the immune environment of the developing brain meninges during synaptogenesis, we performed single cell RNA sequencing (scRNAseq) of immune and non-immune skull-associated meningeal cells on postnatal day (P)14, pre-enriching for lymphocytes and stromal cells (**Fig. 1A; Fig S1A)**. We recovered 12,544 cells for downstream unsupervised clustering analysis (**Fig. 1B, S1C**; quality control in **Fig. S1B**). While cell-type populations were consistent with other studies (DeSisto et al., 2020; Mrdjen et al., 2018; Van Hove et al., 2019; Zelco et al., 2021), the relative abundance of innate to adaptive lymphocytes was markedly different.

**Figure 1:**
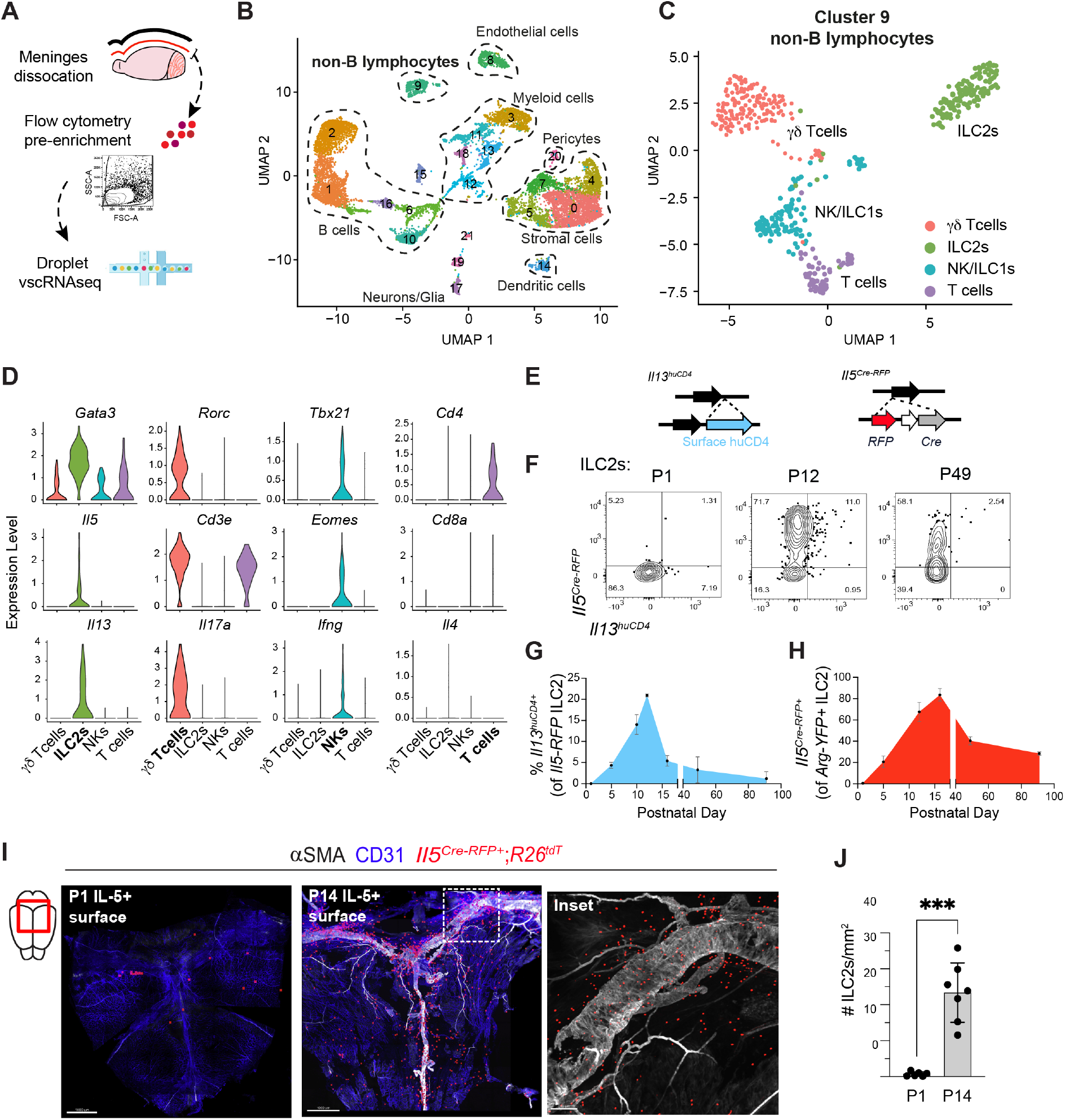
Meningeal type 2 innate lymphocytes expand and produce IL-13 in early life. **A)** Schematic of postnatal day (P)14 dural meninges isolation for flow cytometry and single cell mRNA sequencing. **B)** Unsupervised clustering (UMAP) of 12,544 cells from pooled P14 meninges (enriched for stromal cells and non-myeloid immune cells) with cell-type assignments based on marker gene expression. See Fig. S1A-D for flow cytometry gating, quality control metrics, and cell-type assignment. See Supplemental Table 1 for list of clusterdefining genes. **C)** Sub-clustering of non-B lymphocytes from panel B, including T cells and innate lymphoid cells (Cluster 9; 477 cells). See Supplemental Table 2 for list of cluster-defining genes. **D)** Expression of lineage-defining and effector genes in non-B lymphocytes from panel C. **E)** Schematics depicting cytokine reporter constructs in relation to their respective endogenous murine loci. Note that *Il5^Cre-RFP^* leads to loss of function of the endogenous locus whereas *Il13^huCD4^* is inserted into the 3’ untranslated region and preserves gene expression. *huCD4*: human CD4. **F)** Representative flow plots showing expression of IL-5 (*Il5^Cre-RFP^*) and IL-13 (*Il13^huCD4^*) within Arg1^YFP+^ meningeal ILC2s across development. Gating strategy used to identify ILC2s and specificity of Arg1^YFP^ shown in Fig. S2D. **G)** Quantification of IL-13 expression in meningeal ILC2s over development (percent *Il13^huCD4+^* of *Arg1^YFP+^ Il5^Cre-RFP+^* ILC2s; Mice per age: P1 n=3; P5 n=2; P10 n=3; P12 n=2; P16 n=3; P49 n=3; P91 n=4). **H)** Quantification of meningeal ILC2 IL-5 expression over development (percent *Il5^RFP+^ of Arg1^YFP+^* ILC2s; Mice per age: P1 n=3; P5 n=2; P12 n=2; P16 n=3; P49 n=3; P91 n=3). **I)** Confocal imaging and surface reconstruction of whole-mount dural meninges, highlighting ILC2s (red, *Il5^Cre-RFP^:R26^TdTomato^*), vasculature (blue, CD31) and smooth muscle (white, αSMA) at age P1 and P14. Inset of P14 at the transverse sinus showing native *Il5^Cre-RFP^:R26^TdTomato+^* signal. Scale bar: 1 mm (P1, P14) and 200 μm (inset). **J)** Quantification of meningeal ILC2s (*Il5^Cre-RFP^:R26^TdTomato+^*) at P1 and P14, normalized per area of tissue (P1 n=6, P14 n=7; Dots represent mice; Unpaired t test; ***p < 0.001). Data are mean ± SD.

Innate lymphocytes were predominant at this early life timepoint. Sub-clustering of non-B lymphocytes (Cluster 9) revealed an abundance of ILC2s (*Cd3e^-^, Gata3^+^*) and innate-like gammadelta (*γ*δ) T cells (*Cd3e^+^, Rorc^+^*), as well as smaller populations of putative NK cells (*Cd3e^-^, Tbx21^+^, Eomes^+^*) and group 1 innate lymphoid cells (ILC1s; *Cd3e^-^, Tbx21^+^, Eomes^-^;* **Fig. 1C-D**). In contrast, adaptive immune cells such as CD4^+^ T cells (*Cd3e^+^, Cd4^+^*) were relatively rare. Furthermore, all innate lymphocyte subsets expressed canonical transcription factors and effector genes required for cytokine production, whereas T cells did not (**Fig. 1D, S1D**). We validated these observations at the protein level by flow immunophenotyping over the course of development (**Fig. S2A-C**).

We next quantified effector cytokine production by lymphocytes during development. We found that ILC2s produced a wave of IL-13 and IL-5 activity between postnatal days 5-15, using sensitive transcriptional reporter mice (*Il13^huCD4^:Il5^Cre-RFP^;* **Fig. 1E-H**) and identifying ILC2s with an Arginase-1 reporter (*Arg1^YFP^*, **Fig. S2D-E**; (Schneider et al., 2019)). In contrast, adaptive T cells did not express any detectable IL-5/13 over this same period (**Fig. S2F**). Another type of canonical cytokine, IFNγ (*IFNγ^YFP^*) was tonically expressed in meningeal NK/ILC1 cells (**Fig. S3A-D**). Thus, innate but not adaptive lymphocytes in the brain meninges produce cytokines in early life.

Given this unique burst of ILC2 activation during a critical developmental window and known roles of the type 2 immune axis in tissue remodeling, we further examined meningeal ILC2 expansion during early life. We visualized meningeal ILC2s *in situ* using *Il5^Cre-RFP^;R26^tdTomato^* lineage tracker mice (**Fig. 1I, S2G**). IL-5^+^ cells were rare shortly after birth but expanded to adult levels by P14, localizing to smooth muscle actin-a (aSMA)-rich adventitial regions around the dural sinus and other large vasculature (**Fig. 1I-J**). Together, these results suggest that early-life tissue immune activation is skewed towards type 2 cells (ILC2s) and cytokine release (IL-5/IL-13), suggesting they could be a relevant source of immune signals during postnatal brain development.

### ILC2s increase inhibitory synapse numbers during brain development

During the early postnatal period both excitatory and inhibitory synapses are forming and maturing in the cortex, while ~30% of inhibitory interneurons undergo programmed cell death (Fishell and Kepecs, 2020; Lim et al., 2018; Wong et al., 2018). To determine whether ILC2s impacted these aspects of cortical circuit development we used a well-characterized genetic deletion model to constitutively ablate ILC2s via expression of diphtheria toxin under control of the *Il5* promoter (*Il5^Cre-RFP/Cre-RFP^; R26R*^DTA/DTA^), referred to hereafter as “ΔILC2” (**Fig. 2A**). This deletion strategy is specific to ILC2s in development and targets some subsets of tissue-enriched, innate-like CD4^+^ Th2 cells in later life (Molofsky et al., 2013; Nussbaum et al., 2013; Dahlgren et al., 2019; Cautivo et al., 2022). As the *Il5^Cre-RFP^* construct replaces the endogenous IL-5 locus, all experiments compared ΔILC2 mice to *Il5^Cre-RFP/Cre-RFP^* littermates (“control”). This strategy led to an ~80% depletion of ILC2s without significantly altering other lymphocyte subsets (**Fig. 2B-C**; gating strategy as in **Fig. S2A**).

**Figure 2:**
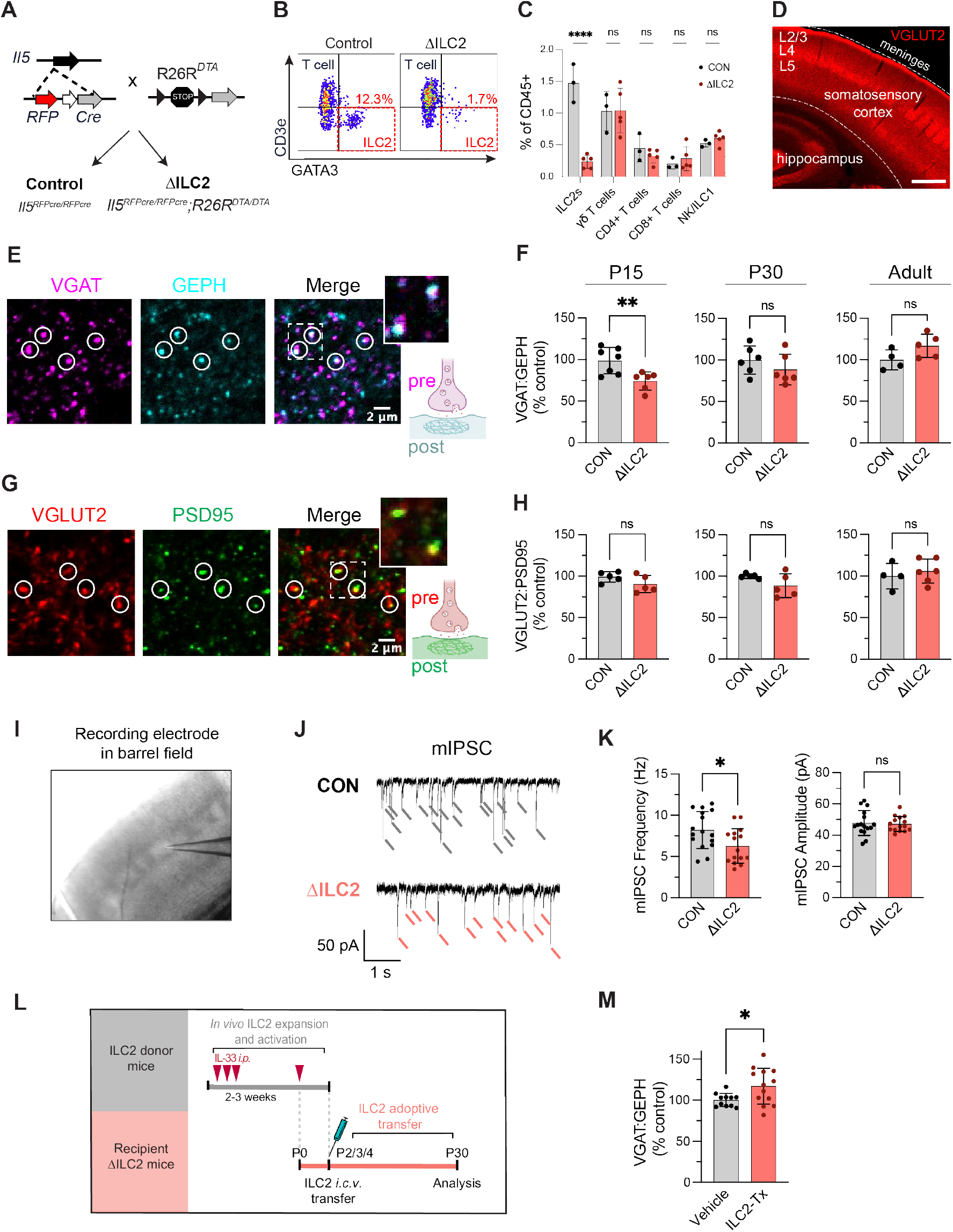
ILC2s increase inhibitory synapse numbers during brain development. **A)** Schematic for generation of ILC2 depleted mice (*Il5^Cre-RFP/Cre-RFP^:R26RP^DTA/DTA^*; a.k.a. “ΔILC2”). All mice were homozygous for *Il5^Cre-RFP^*; ΔILC2 mice were homozygous for *R26R^DTA^*, while controls lacked the DTA allele. Littermates used in all experiments unless otherwise noted. **B)** Representative flow plots showing depletion of meningeal ILC2s (GATA3^+^/CD3^neg^) in ΔILC2 mice compared to controls. Pre-gated on live CD45^+^CD11b^-^CD19^-^ cells (see Fig. S2A for gating). **C)** Quantification of lymphocyte subsets by flow cytometry in ΔILC2 mice as a percentage of controls. Dots represent mice. See Fig. S2A for gating. (Age P15; controls n=3 mice, ΔILC2 n=5 mice; Two-way ANOVA, Šídák’s multiple comparisons test). **D)** Representative coronal section depicting primary somatosensory cortex where synaptic quantifications were performed (VGLUT2 used to highlight cortical layers). **E)** Representative images of inhibitory synapse immunofluorescence staining of colocalized presynaptic (VGAT) and postsynaptic (Gephyrin, GEPH) markers. Circles indicate examples of colocalized puncta; inset area indicated by white dashed box. (Scale bar: 2 μm; inset: 4×4 μm). **F)** Quantification of inhibitory synapses (VGAT:GEPH) in layer 4 somatosensory cortex at the indicated ages. Data at each age is normalized to the mean of the control group within independent experiments (Adult=12-16 weeks; Dots represent mice; Mice per age: P15 n = 7 vs. 6; P30 n = 6 vs. 6; Adult n = 4 vs. 5. Unpaired t-tests). **G)** Representative images of thalamocortical excitatory synapses quantified by colocalization of presynaptic (VGLUT2) and postsynaptic (PSD95^+^) markers. Circles indicate examples of colocalized puncta; inset area indicated by white dashed box (Scale bar: 2 μm; inset: 4×4 μm). **H)** Quantification of excitatory synapses (VGLUT2:PSD95) in layer 4 somatosensory cortex at the indicated ages (Adult=12-16 weeks; Dots represent mice; Mice per age: P15 n = 5 vs. 5; P30 n = 5 vs. 5; Adult n = 4 vs. 6. Unpaired t-tests). **I)** Image of acute cortical slice from whole-cell patch-clamp recordings from excitatory neurons in the somatosensory barrel cortex. **J)** Representative traces of miniature inhibitory postsynaptic current (mIPSC) recordings from acute brain slices of control and ΔILC2 mice. **K)** Quantification of mIPSC frequency (Hz) and amplitude (pA) from the indicated genotypes (Dots represent neurons; n=16 neurons from 4 controls, and 15 neurons from 4 ΔILC2 mice; Age: P28-30; Unpaired t-test.) **L)** Schematic of experimental paradigm for *in vivo* expansion and activation of donor ILC2s and adoptive transfer of ILC2s into ΔILC2 recipient neonatal mice. See Methods for details. **M)** Quantification of inhibitory synapses (VGAT:GEPH) in layer 4 somatosensory cortex at P30 after neonatal transfer of ILC2s (ILC2-Tx). Data is normalized to the mean of the control group within independent experiments. (Dots represent mice; n = 11 vs. 13 per group; 3 independent experiments; Unpaired t-test). Statistics: *p < 0.05, **p < 0.01, ****p < 0.0001. Data are mean ± SD.

We first assessed whether ΔILC2 mice had altered numbers of excitatory and inhibitory synapses. We focused on somatosensory cortex, where key landmarks enable unbiased and reproducible quantification (S1**, Fig. 2D**). We observed a significant decrease in inhibitory synapses in ΔILC2 mice at P15 as measured by colocalization of the pre- and post-synaptic markers VGAT and Gephyrin (**Fig. 2E-F;** quantification in cortical layer 4 (L4)). This decrease was most prominent at P15 and no longer present by adulthood (**Fig. 2F**). In contrast, excitatory thalamocortical synapses (VGLUT2^+^PSD95^+^) in this region were unaltered (**Fig. 2G-H**). Whole cell patch clamp recordings from excitatory neurons in L4 barrels revealed a reduction in miniature inhibitory post-synaptic currents (mIPSCs), consistent with a reduction in synapse numbers, whereas amplitude and kinetics were unchanged (**Fig. 2I-K, Fig. S4G**).

Inhibitory synapses represent inputs from a diverse group of inhibitory neuron subsets. We separately characterized axo-somatic synapses, which are formed primarily by parvalbumin^+^ basket cells, and axo-axonic synapses onto the axon initial segment, which are made by chandelier cells (Tremblay et al., 2016). Neither of these were altered in ΔILC2 mice (**Fig. S4A-E**). These data may suggest a preferential effect of ILC2s on axo-dendritic synapses, which are the predominant synapse type in the developing cortex. Of these, we observed a similar reduction across both superficial (L2/3) and deeper (L4) cortical layers (**Fig. S4F**). We did not observe changes in inhibitory neuron numbers or distribution across cortical layers as assessed by GAD67^+^, PV^+^, and SST^+^ soma at P15, a time point when developmental apoptosis is largely complete and interneuron numbers have stabilized (Lim et al., 2018; Wong et al., 2018); **Fig. S4H-I**). These data indicate that ILC2s preferentially impact axo-dendritic inhibitory synapse formation.

To determine whether ILC2s were sufficient to increase inhibitory synapses during this period, we transferred activated ILC2s into neonatal ΔILC2 mice. ILC2s were stimulated to produce cytokine *in vivo* by IL-33 treatment of donor mice (Cautivo et al., 2022). ILC2s isolated from lung were delivered by intracerebroventricular (i.c.v.) injection into ΔILC2 recipients between P2-4 (**Fig. 2L**). Mice that received activated ILC2s had significantly increased inhibitory synapses compared to vehicle-injected littermates (**Fig. 2M;** P30). Taken together, these data suggest that ILC2s promote inhibitory synapse formation during a postnatal developmental window that closely follows their peak of IL-13 production.

### IL-13/4 signaling to interneurons increases inhibitory synapse numbers during brain development

To directly assess whether IL-13 could impact inhibitory synapse numbers, we quantified cortical synapses in both loss- and gain-of-function models. IL-13 signals via a heterodimeric receptor consisting of the IL-13Rα1 and IL-4Rα subunits (**Fig. 3A**; (Van Dyken and Locksley, 2013)). Genetic loss of either obligate subunit blocks IL-13 signaling. We found that global deficiency of the *Il4ra* subunit (*Il4ra^-/-^*) phenocopied the reduction of inhibitory synapses seen in ILC2-deficient animals at P15 and did not impact excitatory synapse numbers (**Fig. 3B**). Conversely, direct delivery of IL-13 by i.c.v. injection was sufficient to increase inhibitory synapses within 20 hours and did not impact excitatory synapse numbers (**Fig. 3C**). These data demonstrate that IL-13/4 signaling is sufficient to increase inhibitory synapse numbers.

**Figure 3:**
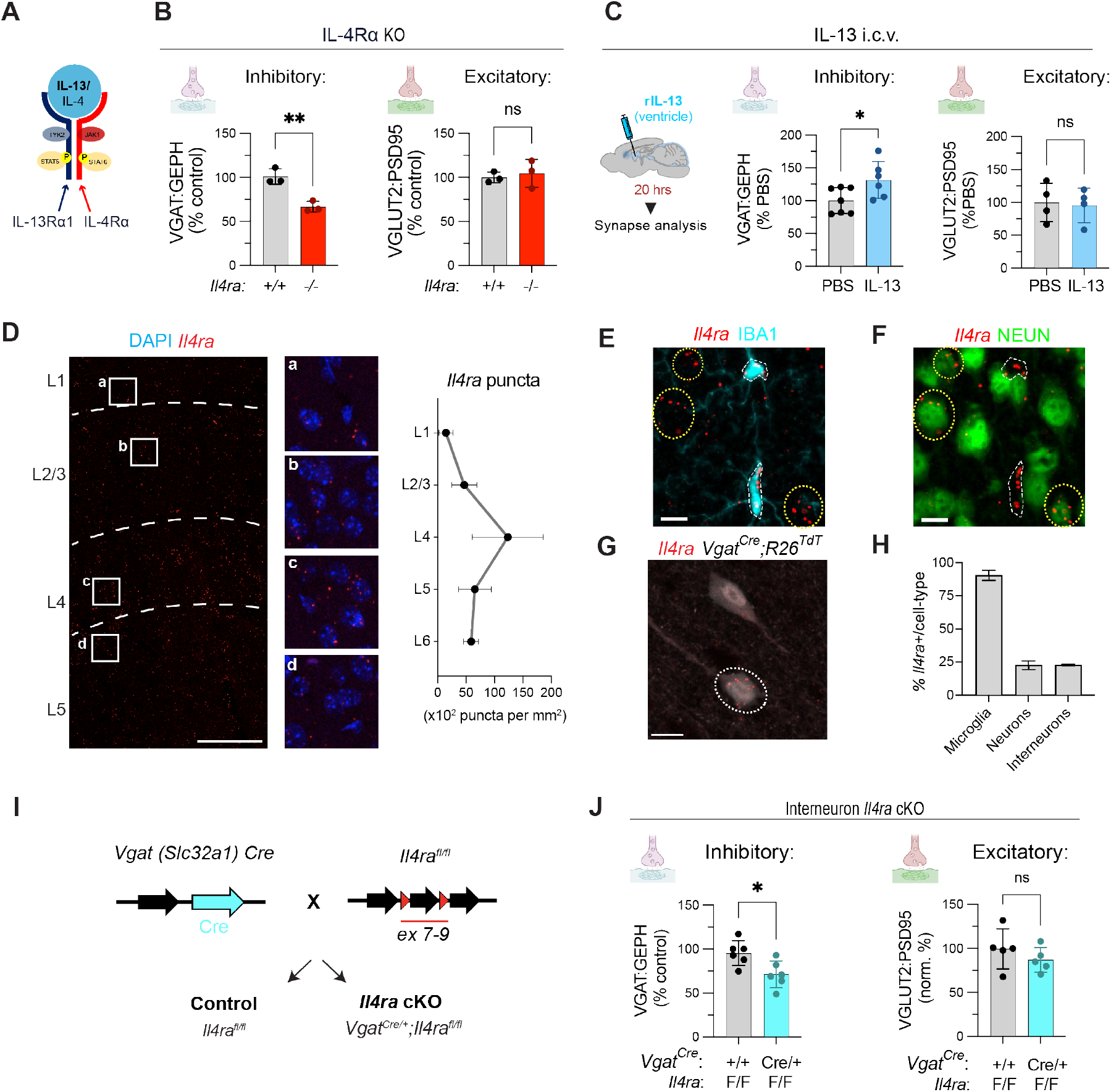
IL-13/4 signaling to interneurons increase inhibitory synapse numbers during brain development. **A)** Schematic of the IL-13/4 receptor consisting of the IL-13Rα1 and IL-4Rα subunits. **B)** Quantification of inhibitory (VGAT^+^GEPH^+^) and excitatory (VGLUT2^+^PSD95^+^) synapses in *Il4ra^+/+^* vs. *Il4ra^-/-^* mice (Age P15; n=3 mice per group; Unpaired t test). **C)** Quantification of inhibitory and excitatory synapses 20 hours post-i.c.v. injection of IL-13 (250 ng) or PBS (Age P15; Inhibitory: n=7 control, n=6 IL-13-injected mice; Excitatory: n=4 mice per group; 3 independent experiments; Unpaired t test). **D)** Quantitative *in situ* hybridization for *Il4ra* counterstained with DAPI in the somatosensory cortex. Insets (a-d) highlight cortical layers (Age P30; 20X tiled image; scale bar: 100 μm). **Right**: Quantification of *Il4ra* puncta density by layer (n=3 mice; Age P30-60; Dots represent mean). **E)** Representative image showing colocalization of *Il4ra* transcript with microglia (IBA1^+^; scale bar: 10 μm). **F)** Representative image showing colocalization of *Il4ra* transcript with neurons (NEUN^+^; scale bar: 10 μm). **G)** Representative image showing colocalization of *Il4ra* transcript with an inhibitory interneuron reporter (*Vgat^Cre/+^;R26^tdTomato+^;* scale bar: 10 μm). **H)** Quantification of percentage of *Il4ra^+^* cells per cell-type in somatosensory cortex, as in E-G; n=2 mice). **I)** Genetic constructs used to conditionally delete *Il4ra* from inhibitory neurons (“interneuron *Il4ra* cKO”). Cre is inserted after the stop codon of the endogenous *Slc32a1 (Vgat*) locus (Vong et al. 2011). The *Il4ra^flox^* allele allows Cre-mediated excision and conditional deletion of exons 7-9 of *Il4ra* (Herbert et al 2004). **J)** Inhibitory (VGAT^+^GEPH^+^) and excitatory (VGLUT2^+^PSD95^+^) synapse quantifications in interneuron *Il4ra* cKO vs. control mice (Age P30; Inhibitory, n=6 mice/genotype; Excitatory, n=5 mice/genotype; 2 independent experiments; Unpaired t tests).

To determine the cellular targets of IL-13 signaling in the developing cortex, we examined expression of the IL-13 receptor subunit *Il4ra. Il4ra* mRNA was detected throughout cortical layers with enrichment in L4 (**Fig. 3D**). Co-immunostaining revealed that almost all microglial and myeloid cells expressed *Il4ra* (>90% of IBA1^+^ cells, **Fig. 3E,H**), as did subsets of NEUN^+^ neurons (~25% of NEUN^+^ cells, **Fig. 3F,H**) and *Vgat^Cre/+^;R26^tdTomato+^* interneurons (~25% of *tdT*^+^ cells, **Fig. 3G,H**), in line with published datasets from murine cortex (Yao et al., 2021). Single cell RNAseq of non-neuronal cells in the developing cortex (Dorman et al., 2022) and our meninges scRNAseq dataset confirmed expression of both receptor subunits in myeloid cells, including microglia and meningeal macrophages (**Fig. S5A-B**). Therefore, both neurons and myeloid cells are relevant cellular targets of IL-13 signaling during development.

To determine if microglia or macrophages might mediate the impact of IL-13 on inhibitory synapses, and given their role in inhibitory synaptic remodeling (Favuzzi et al., 2021), we conditionally deleted *Il4ra* in both microglia and all myeloid cells using a Cre-dependent *Il4ra^flox^* model (Herbert et al., 2004). While microglia expressed cell surface IL-4Rα and had a robust transcriptional response to exogenous IL-13 *in vivo* **(Fig. S5C-D**), there was no impact on inhibitory synapse numbers following conditional deletion of the IL-13/4 receptor in microglia (*P2ry12^CreERT^*, **Fig. S5E-F**) or in all macrophages (*Cx3cr1^CreERT^*, **Fig. S5G-J**;) (Mckinsey et al., 2020; Yona et al., 2013). Thus, IL-13 signaling to myeloid cells is dispensable for inhibitory synapse formation.

In contrast, we observed a significant decrease in inhibitory synapses when we conditionally deleted *Il4ra* from GABAergic inhibitory neurons (**Fig. 3I-J***; Vgat^Cre/+^ :Il4ra^flox/flox^* “interneuron *Il4ra* cKO”, (Vong et al., 2011)). Excitatory synapse numbers were not altered in these mice (**Fig. 3J**). As previously reported, the presence of the *Cre* allele on its own had no effect on synapse numbers (**Fig. S5K**). Deletion of *Il4ra* with a pan-neuronal transgenic Cre (*Syn1^Cre+^; Il4ra^flox/flox^*; (Zhu et al., 2001)) showed a similar trend (**Fig. S5L**). Thus, IL-13 signaling directly to inhibitory neurons promotes inhibitory synapse formation and phenocopies the synaptic impact of ILC2 deletion.

### ILC2s and IL-13 signaling to interneurons promote social interaction in adulthood

Synaptic inhibition plays critical roles in tuning overall network activity (Marín, 2012), and transient perturbations in circuit maturation during development impact adult social behavior (Bitzenhofer et al., 2021; Magno et al., 2021; Reed et al., 2019). To determine how this developmental neuroimmune circuit influenced adult behavior, we examined social, object, and contextual fear memory in ΔILC2 mice and littermate controls (age 2-3 months). In the three-chamber social interaction test (**Fig. 4A**), mice were first habituated to the chamber then allowed to explore the field containing a novel mouse placed under a wire cup in one chamber and an empty wire cup in the other. While controls preferred the social cup to the empty cup, ΔILC2 showed reduced social interaction and a significant decrease in discrimination index relative to controls, indicating impaired sociability (**Fig. 4B-D**). Preference for social novelty was not significantly altered, as assessed 24 hours later by placing test mice in the three-chamber apparatus containing the familiar mouse from day 1 and a novel mouse (**S6A-B**). Other behavioral metrics were not altered, including: object memory in the novel object recognition (NOR) assay; mobility in an open field; anxiety-like behavior in the elevated plus maze; and learning, spatial memory, and context discrimination in a contextual fear conditioning assay (**Fig. S6C-I**). Together, these data suggest a preferential impact of ILC2s on adult social behavior.

**Figure 4:**
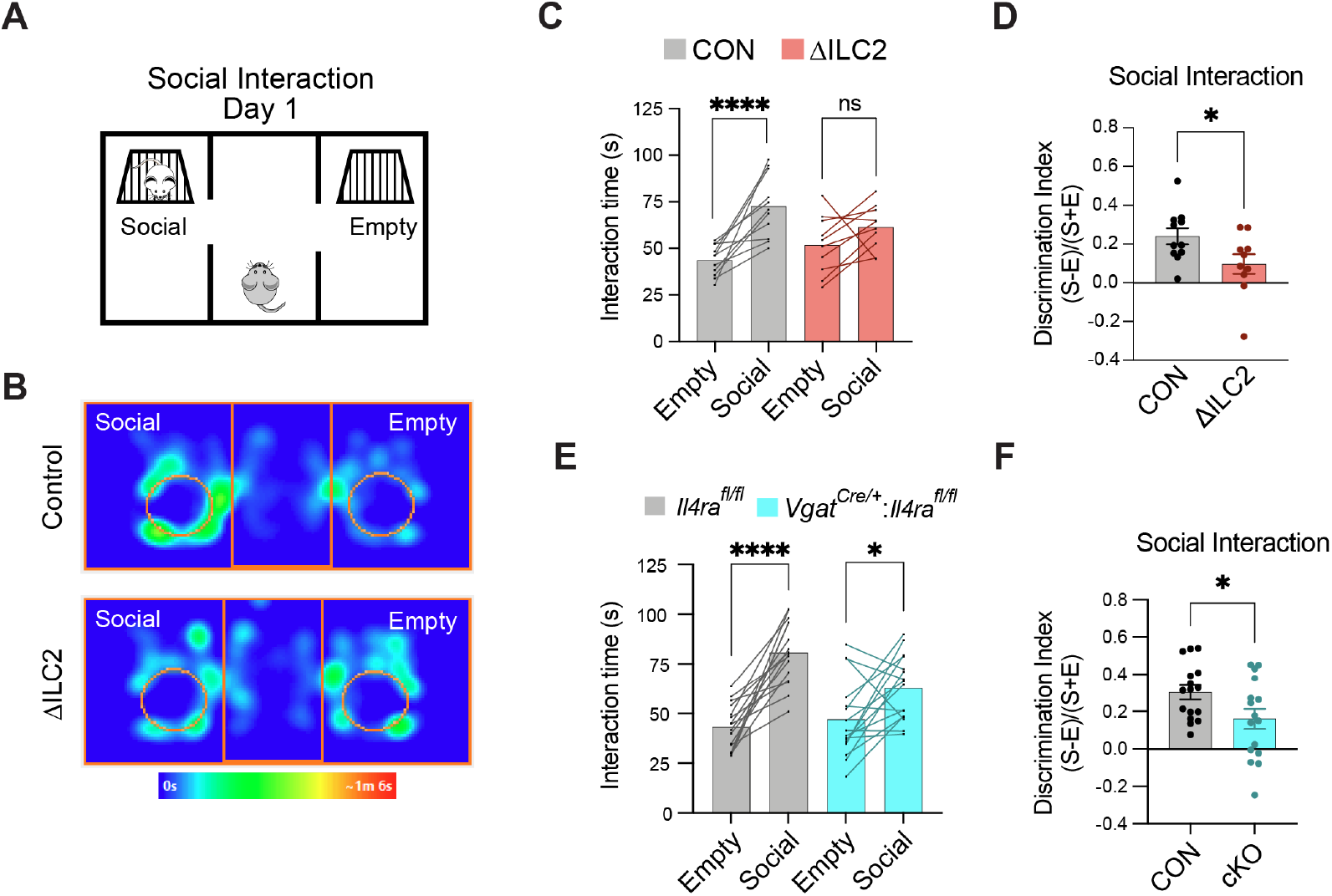
ILC2s and IL-13 signaling to interneurons promote social interaction in adulthood. **A)** Schematic of Crawley’s three-chamber assay of social interaction (Day 1 of social testing). **B)** Representative mouse tracking heatmaps of time spent interacting with social or empty stimuli by control mice vs. ΔILC2 mice, revealing impaired social preference. **C)** Interaction time spent with the Empty vs. Social cup on Day 1 for control mice (*Il5^Cre-RFP/Cre-RFP^*) vs. ΔILC2 mice (*Il5^Cre-RFP/Cre-RFP^:R26R*^DTA/DTA^; Two-way ANOVA, Šídák’s multiple comparisons test). n = 11 vs. 10 mice/genotype. Data are means with lines connecting paired data for each mouse. ****p < 0.0001. **D)** The social interaction Discrimination Index for ΔILC2 mice and controls calculated as the difference in interaction time with Social (S) and Empty (E) cup, over total interaction time ((S-E)/(S+E); Unpaired t test). Dots=mice. mean ± SEM. *p < 0.05. **E)** Interaction time spent with the Empty vs. Social cup on Day 1 of the social interaction assay by controls vs. interneuron *Il4ra* cKO mice (Two-way ANOVA, Šídák’s multiple comparisons test). n=16 mice/genotype. Data are means with lines connecting paired data for each mouse. *p < 0.05, ****p < 0.0001. **F)** The social interaction discrimination Index for cKO mice and controls calculated as the difference in interaction time with social (S) vs. empty (E) cup, over total interaction time, for control vs. interneuron *Il4ra* cKO mice ((S-E)/(S+E); Unpaired t test). Dots=mice. mean ± SEM. *p < 0.05.

Given the phenocopy of inhibitory synaptic defects in interneuron *Il4ra* cKO animals, we also performed behavioral assays in these animals. Like ILC2-deficient mice, interneuron *Il4ra* cKO animals had significantly reduced social interaction in the three-chamber assay relative to littermate controls (**Fig. 4E-F**). Social novelty preference on day two of testing was not altered, nor were other behavioral measures including open field and contextual fear conditioning (**Fig. S6K-L**). Thus, both ILC2 deficiency and loss of IL-13 signaling to interneurons preferentially impacted social behavior. Overall, these data suggest a model whereby ILC2-derived IL-13 increases inhibitory synapses during development and promotes social behavior in adulthood (**Fig. 5**).

**Figure 5:**
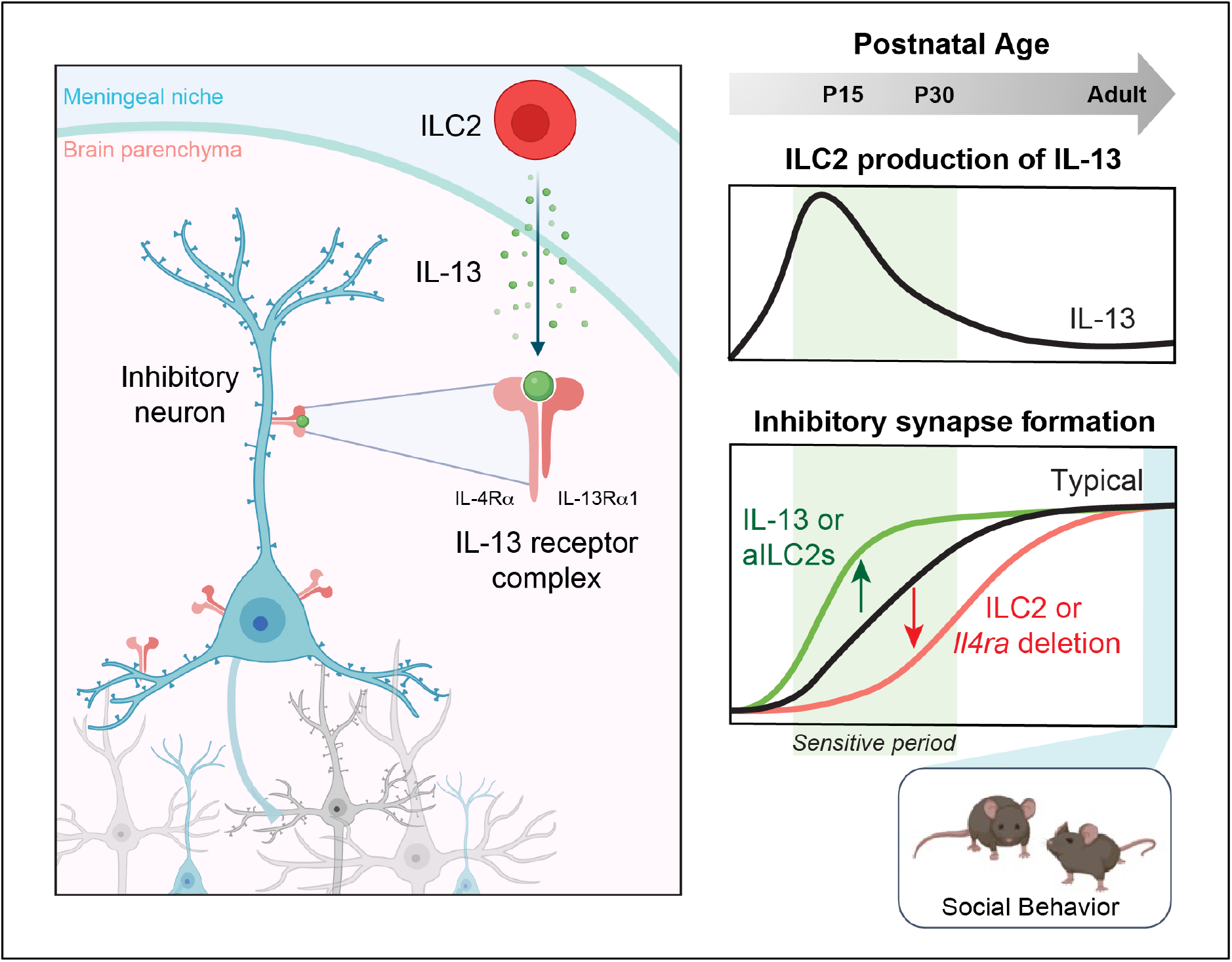
Graphical Abstract. **A)** Group 2 innate lymphoid cells (ILC2s) promote early-life inhibitory synapse development and social behavior. This study supports a model in which Interleukin-13 (IL-13) derived from activated, early-life ILC2s signals directly to GABAergic cortical neurons to promote inhibitory synapse maturation. More broadly, they suggest that innate immune signaling can dynamically shape neuronal connectivity during key periods of brain development. (aILC2s, activated ILC2s).

## DISCUSSION

Despite an emerging recognition of innate lymphocytes as central players in tissue immunity, it is not clear that they are essential for host defense, raising the question of their role in homeostasis (Kotas and Locksley, 2018). We observed an expansion of innate lymphocytes in the early postnatal brain meninges. ILC2s were the dominant source of IL-13 in the meninges and produced a surge of IL-13 between 5-15 postnatal days, mirroring a similar wave of ILC2 expansion and activation in other tissues (Ricardo-Gonzalez et al., 2018; Schneider et al., 2019) In contrast, adaptive lymphocytes were quiescent during development and expanded by adulthood. In the setting of the neonatal lung, ILC2s function to promote and maintain a type 2 immune environment in homeostasis (Saluzzo et al., 2017). It is interesting to speculate that one role of this type 2 developmental wave is to synchronize development across organ systems, including across brain and body.

This work reveals a novel role for ILC2s and IL-13 signaling to interneurons to promote inhibitory synapses in early life and sociability in adulthood. Furthermore, we demonstrate a unique developmental role for IL-13 signaling to interneurons that may be distinct from other roles for IL-4 receptor in the adult brain, which are ILC2-independent (Derecki et al., 2010; Herz et al., 2021; Vogelaar et al., 2018; Hanuscheck et al., 2022; Brombacher et al., 2017, 2020; Li et al., 2023). While these and other studies have clearly demonstrated that lymphocyte-derived cytokines impact the brain, these cells are largely outside the blood brain barrier (BBB). Emerging evidence suggests that the BBB is dynamic and permeable to biologically active molecules (Yang et al., 2020). Defining the permeability of the BBB during development, and the routes through which cytokines reach the CNS parenchyma is an essential future direction in understanding the brain-immune interface.

Our study shows that IL-13 signals directly to inhibitory interneurons to promote inhibitory synapse formation. Interneurons represent a minority (~20%) of cortical neurons but can dynamically modulate brain function (Fishell and Kepecs, 2020). Alterations in inhibition can have broad impacts, from preventing hyperexcitability and seizures, to more complex effects on adult social behavior and cognition (Rubenstein, 2010; Sohal and Rubenstein, 2019). Our study is consistent with others demonstrating that cytokine receptor expression on neurons impacts brain development and social behavior (Choi et al., 2016; Reed et al., 2019). It is also consistent with studies demonstrating direct cytokine signaling to interneurons (Filiano et al., 2016; Herz et al., 2021). Given that interneurons are morphologically, molecularly, and functionally diverse (Tremblay et al., 2016), determining whether interneuron subtypes are differentially responsive to the ILC2/IL-13 axis is a key future direction.

Ultimately, these findings raise the question of whether type 2 immune signals might impact human social or cognitive development. Alterations in excitatory/inhibitory synaptic balance are implicated in disorders including epilepsy, autism spectrum disorders, and schizophrenia. Epidemiologic links to immune dysfunction in these conditions have repeatedly been observed (Bennett and Molofsky, 2019). Type 2 immunomodulators that could rapidly alter inhibitory tone in the brain might represent a powerful therapeutic strategy in these disorders. Conversely, these data raise the question of how environmental stressors that alter type 2 immune tone might impact cognitive or social development. High parasite burden is a common feature of childhood in many parts of the world and is frequently associated with delays in learning and cognition (Fernando et al., 2010). Although the psychosocial and metabolic contributions to this effect are complex, directly examining how allergic immune challenges impact brain development could have major impacts on human health.

## Supplemental Figures

**Figure S1:**
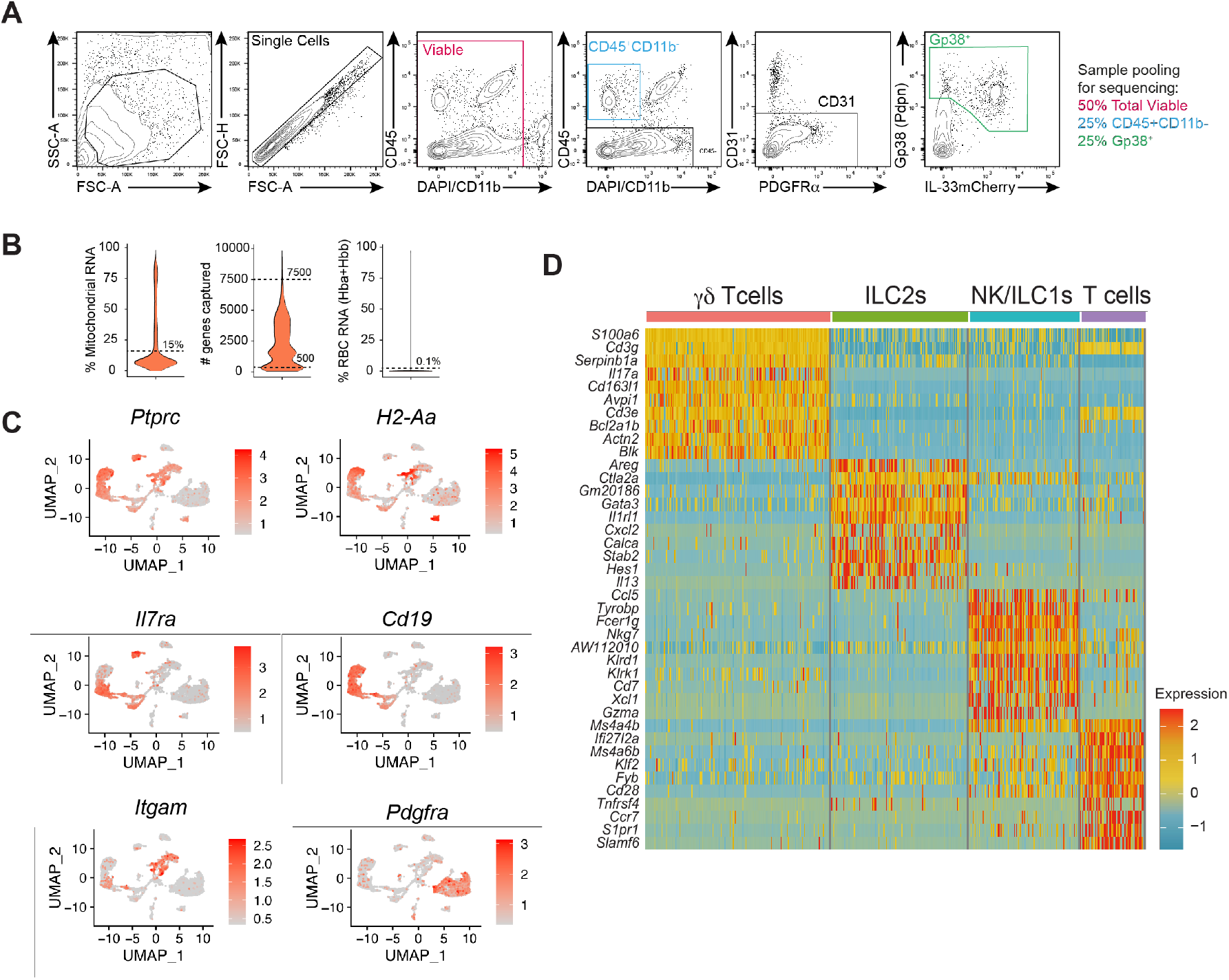
Gating and quality control for meningeal single cell profiling. **A)** Gating strategy for fluorescence activated cell sorting of P14 dural meninges for scRNAseq in Fig. 1, showing gates used to enrich non-myeloid immune cells (blue, CD45^+^CD11b^-^, 25% of final pool) and stromal cells (green, gp38^+^CD31^-^, 25% of final pool) and total viable (red, 50% of final pool). Genotypes of mice included were *Il33^mcherry/+^* and *Il33^mcherry/mcherry^* however differences were modest and were not analyzed further. **B)** Quality control metrics for single cell sequencing, thresholds indicated with dotted lines. See Methods for details. **C)** Feature plots of data in Fig. 1B, showing representative genes used for cell-type assignment. *Ptprc:* hematopoietic cells; *H2-Aa:* MHC class II^+^ myeloid subsets; *Il7ra:* lymphocytes; *Cd19:* B lymphocytes; *Itgam:* macrophage subsets; *Pdgfra:* stromal cells. **D)** Heatmap of top cluster-defining genes by log fold change for non-B lymphocytes in Fig. 1C used for cell-type assignments.

**Figure S2:**
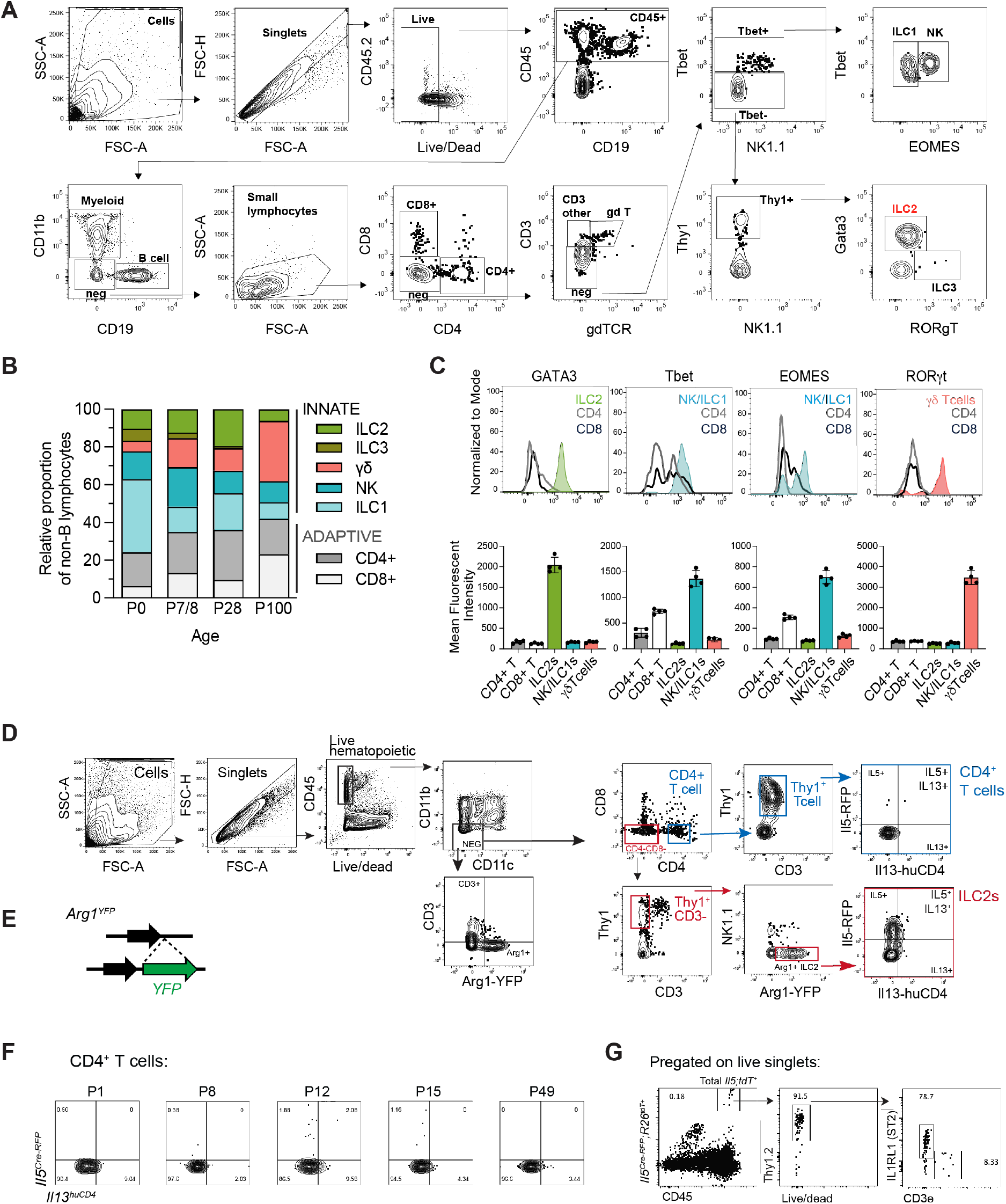
Gating strategies and additional cytokine reporter comparisons. **A)** Flow cytometry gating strategy for analysis of meningeal lymphocyte subsets. **B)** Quantification by flow cytometry of innate and adaptive lymphocytes in meninges across development (pre-gated on viable CD45^+^CD11b^-^CD19^-^ cells, shown as a relative proportion among the indicated cell types; Mice per age: P0 n=2; P7/8 n=4; P28 n=3; P100 n=4; Gating strategy in A). **C)** Representative flow histograms (top) and quantification (bottom) of protein expression of lineage-defining transcription factors including GATA3 (type 2), Tbet (type 1), EOMES (type 1), and RORgt (type 3/17). (P28 n=4; Dots represent mice. Gating strategy in S1H). **D)** Flow cytometric gating strategy for identification of meningeal ILC2s and CD4 T cells in *Arg1^YFP/YFP^; Il5^Cre-RFP/+^; Il13^huCD4/+^* mice. ILC2 gating strategy in red, CD4 T cell gating strategy in blue. **E)** Schematic depicting *Arg1^YFP^* cytokine reporter construct inserted in the endogenous 3’UTR of the murine *Arg1* locus which preserves gene expression. **F)** Representative flow plots showing expression of IL-5 (*Il5^Cre-RFP^*) and IL-13 (*Il13^huCD4^*) in meningeal Thy1^+^ CD4^+^ T cells across development. **G)** Representative flow plots from meninges of *Il5^Cre-RFP^:R26^TdTomato^* lineage tracker mice, demonstrating a majority of *Il5^Cre-RFP^:R26^TdTomato+^* hematopoietic cells are ILC2s (e.g. Thy1^+^ IL1RL1/ST2^+^).

**Figure S3:**
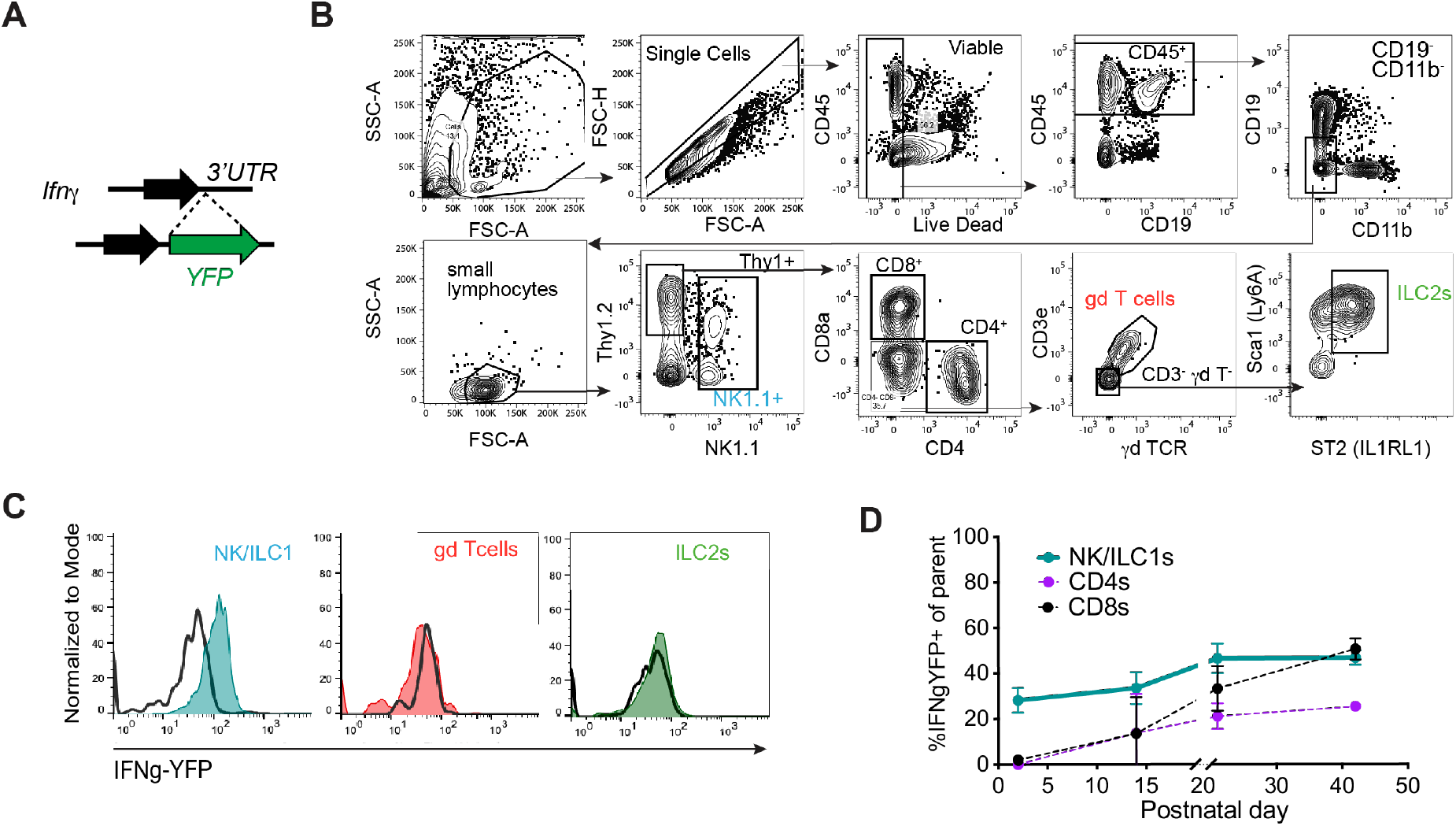
*Ifng^YFP^* cytokine reporter gating strategy and meningeal analysis. **A)** Schematic depicting *Ifng^YFP^* genetic construct inserted in the 3’UTR of the murine *Ifng* locus. **B)** Gating strategy of meningeal lymphocytes in *Ifng^YFP^* mice (NK1.1^+^ ILC1s/NK cells, CD8^+^ T cells, CD4^+^ T cells, CD3e^+^ TCRγδ^+^ γδ T cells, and lineage^-^ Thy1^+^ ST2(IL1RL1)^+^ Sca-1^+^ ILC2s). **C)** Representative flow histograms depicting IFNγ (*Ifng^YFP^*) expression in meningeal NK/ILC1s (teal), γδ T cells (red), and ILC2s (green) compared to YFP^-^ WT mice (black line). **D)** Quantification of IFNγ (*Ifng^YFP^*) expression in meningeal NK/ILC1s, CD8^+^ T cells, and CD4^+^ T cells over development, defined as percent of cell type positive for *Ifng^YFP^*. (n=4 mice/age, except P42 n=3). Data are mean ± SD.

**Figure S4:**
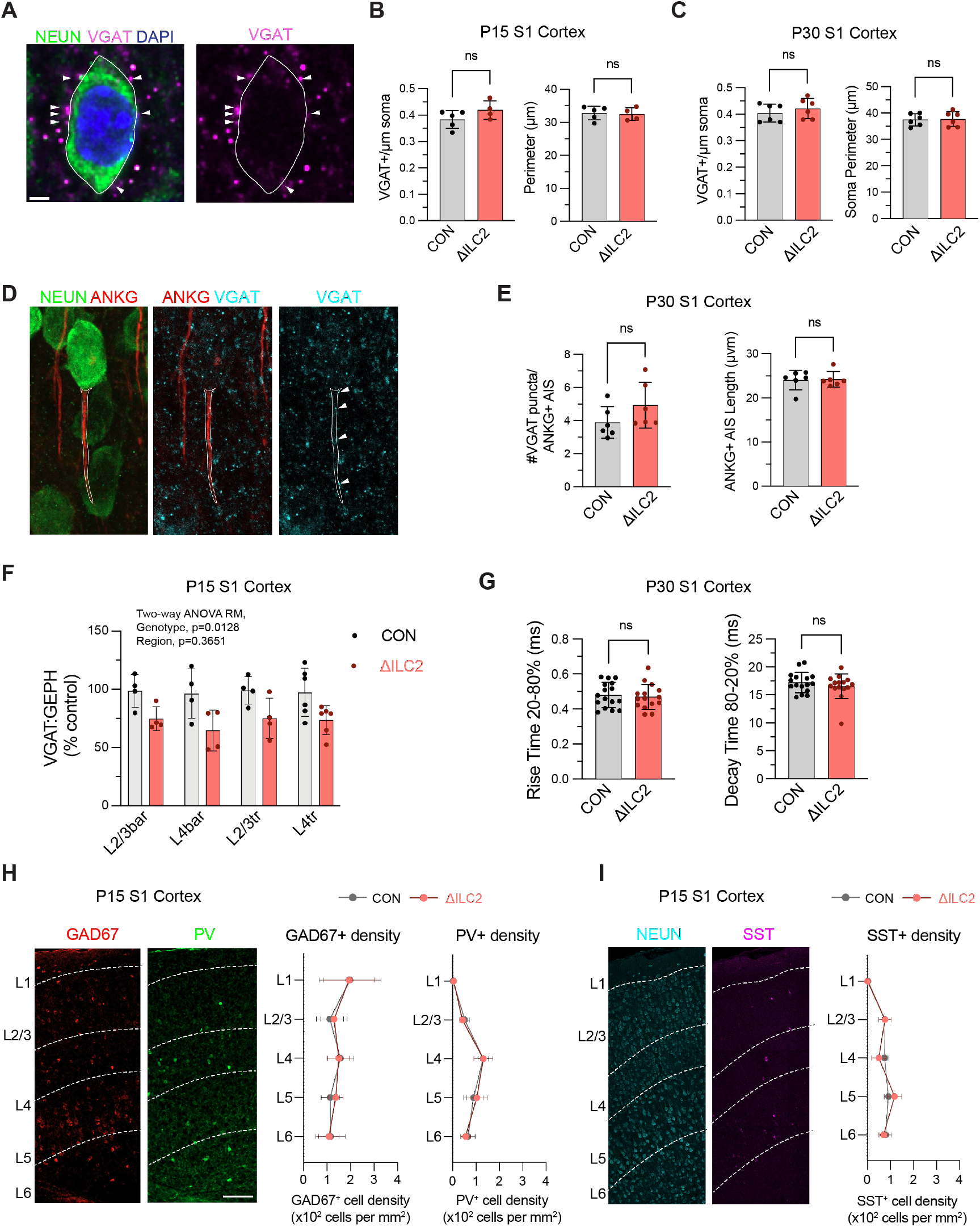
Impact of ILC2s on inhibitory synapses and interneuron numbers. **A)** Representative immunostaining for neuronal soma (NEUN) and the presynaptic marker VGAT in the somatosensory cortex. White arrowheads indicate perisomatic synapses. Scale bar: 2 μm. **B)** Quantification of perisomatic synapse density (VGAT puncta/μm soma) and soma perimeter (μm) at P15. (Dots represent mice; n=5 vs. 4; 40-45 individual neurons; Unpaired t tests). **C)** Quantification of perisomatic synapse density (VGAT puncta/μm soma) and soma perimeter (μm) at P30. (Dots represent mice; n=6 vs. 6; 40-41 individual neurons; Unpaired t tests). **D)** Representative immunostaining for neuronal soma (NEUN), axon initial segment (AIS) protein Ankyrin G (ANKG), and the presynaptic marker (VGAT) in the somatosensory cortex. **E)** Quantification of AIS synapse density (VGAT puncta/AIS) and AIS length (μm) at P30. (Dots represent mice; n=6 vs. 6; 153-167 individual AIS; Unpaired t tests). **F)** Quantification of inhibitory synapses by colocalization of presynaptic (VGAT) and postsynaptic (Gephyrin) markers in multiple S1 layers (L2/3 and L4) and regions (*bar*, barrel field; *tr*, trunk field). Data normalized to mean of controls within independent experiments. (Age P15; Dots represent mice; n=4-6 mice/genotype, 2-3 independent experiments; Two-way RM ANOVA: Effect by genotype: p=0.0128, Effect by region: p=0.3651). **G)** Additional mIPSC parameters from patching experiments in Fig. 2K. (Dots represent neurons; n=16 neurons from 4 controls, and 15 neurons from 4 ΔILC2 mice; Age P28-30; Unpaired t-test.) **H)** Left: Representative staining of GAD67 and Parvalbumin (PV) in P15 S1 cortex. Scale bar: 100 μm. Right: Quantification by cortical layer of total inhibitory neurons (GAD67^+^) and Parvalbumin (PV^+^) neurons in somatosensory cortex (Age P15; n=5 controls, 4 ΔILC2 mice; 2 independent experiments; Two-way ANOVA, Šídák’s multiple comparisons test; Data are mean ± SD). **I)** Left: Representative staining of NEUN and somatostatin (SST) in P15 S1 cortex. Right: Quantification by cortical layer of SST^+^ neurons in somatosensory cortex (Age P15; n=5 controls, 5 ΔILC2 mice; 2 independent experiments; Two-way ANOVA, Šídák’s multiple comparisons test; Data are mean ± SD).

**Figure S5:**
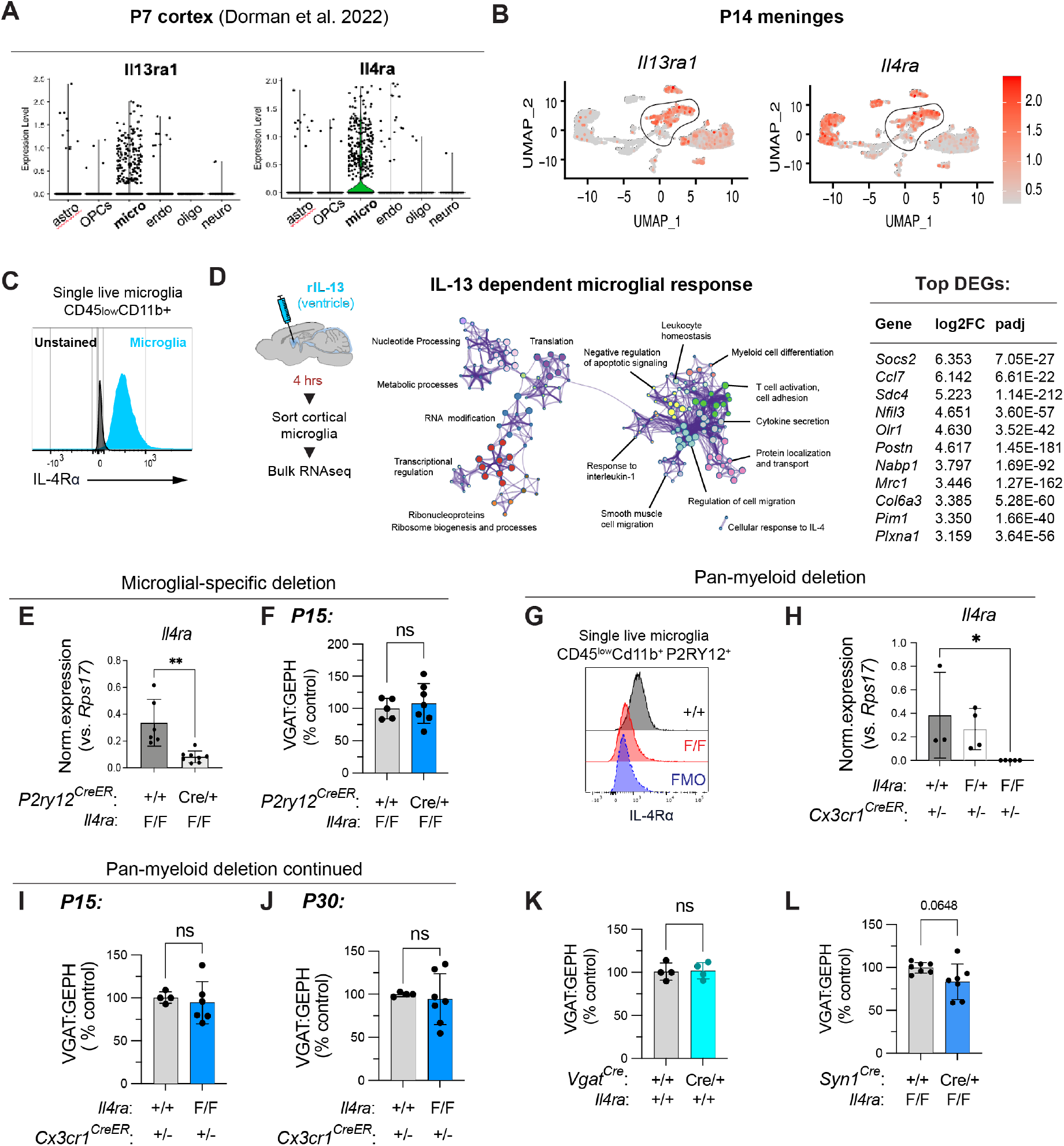
Myeloid cells respond to IL-13 but are dispensable for inhibitory synapse formation. **A)** Expression of *Il13ra1* and *Il4ra* in single cell RNA sequencing of primarily non-neuronal brain cells at age P7 (generated from (Dorman et al., 2022) dataset). **B)** Expression of *Il13ra1* and *Il4ra* in single cell RNA sequencing of meningeal cells at age P14 from data in Fig. 1. **C)** Representative histogram of flow cytometry data showing protein expression of IL-4Rα in microglia isolated from P15 mice. Microglia (blue, CD45^low^ CD11b^+^) compared to unstained cells (black). **D)** Schematic of bulk RNA sequencing experiment after intracerebroventricular injection of IL-13 (250 ng) or PBS in P30 mice (n=4 mice/group). Cortical microglia were FACS sorted 4h post-injection for RNAseq library prep. Center shows Metascape GO term enrichment analysis of differentially upregulated genes (DEGs); Top DEGs shown to right. **E)** Quantitative PCR on sorted microglia after conditional deletion of IL-4Rα in microglia (*P2ry12^CreERT/+^;Il4ra^flox/flox^* and *Il4ra^flox/flox^* mice treated neonatally with tamoxifen; see Methods). *Il4ra* expression normalized to the housekeeper gene *Rps17* (Age P15; n=6 controls, 8 cKOs; 2 independent experiments; Unpaired t test). **F)** Inhibitory synapse quantification in microglia-specific *Il4ra* cKO mice (*P2ry12^CreERT/+^;Il4ra^flox/flox^*) vs. controls (*Il4ra^flox/flox^;* Age P15; n=5 controls, 7 cKOs; 3 independent experiments; Unpaired t test). **G)** Representative flow cytometry histogram of IL-4Rα protein on microglia after conditional *Il4ra* deletion (pre-gated for CD45^+^CD11b^+^P2RY12^+^ cells). +/+ control (black, *Cx3cr1^CreERT/+^;Il4ra^+/+^*), F/F cKO (red, *Cx3cr1^CreERT/+^;Il4ra^flox/flox^*), FMO (blue, fluorescence minus one for IL-4Rα stain on control microglia). **H)** Quantitative PCR on sorted microglia after conditional deletion of IL-4Rα in myeloid cells (*Cx3cr1^CreERT/+^;Il4ra^flox/flox^, Cx3cr1^CreERT/+^;Il4ra^flox/+^*, and *Cx3cr1^CreERT/+^;Il4ra^+/+^* mice treated neonatally with tamoxifen; see Methods). *Il4ra* expression normalized to the housekeeper gene *Rps17* (Age P15; n=3 controls, 4 heterozygotes, and 5 cKOS; 3 independent experiments. One-way ANOVA). **I)** Inhibitory synapse quantification in pan-myeloid *Il4ra* cKO mice (*Cx3cr1^CreERT/+^;Il4ra^flox/flox^*) vs. controls (*Cx3cr1^CreERT/+^;Il4ra^+/+^*) at P15 (n=4 controls, 6 cKOs. 3 independent experiments; Unpaired t test). **J)** Inhibitory synapse quantification in pan-myeloid *Il4ra* cKO mice (*Cx3cr1^CreERT/+^;Il4ra^flox/flox^*) vs. controls (*Cx3cr1^CreERT/+^;Il4ra^+/+^*) at P30 (n=4 controls, 7 cKOs. 3 independent experiments; Unpaired t test). **K)** Control experiment to assess the impact of *Vgat^Cre^* allele on inhibitory synapse numbers (comparison of *Vgat^Cre/+^* vs. wildtype, in the absence of a floxed allele; Age P30; n=4 mice/group; 3 independent experiments; Unpaired t test). **L)** Inhibitory synapse quantification in *Syn1^Cre/+^:Il4ra^flox/flox^* pan-neuronal cKO vs. *Il4ra^flox/flox^* control mice (Age P30; n=7 mice/genotype; 3 independent experiments; Unpaired t test). Data are mean ± SD. Statistics: *p < 0.05, **p < 0.01.

**Figure S6:**
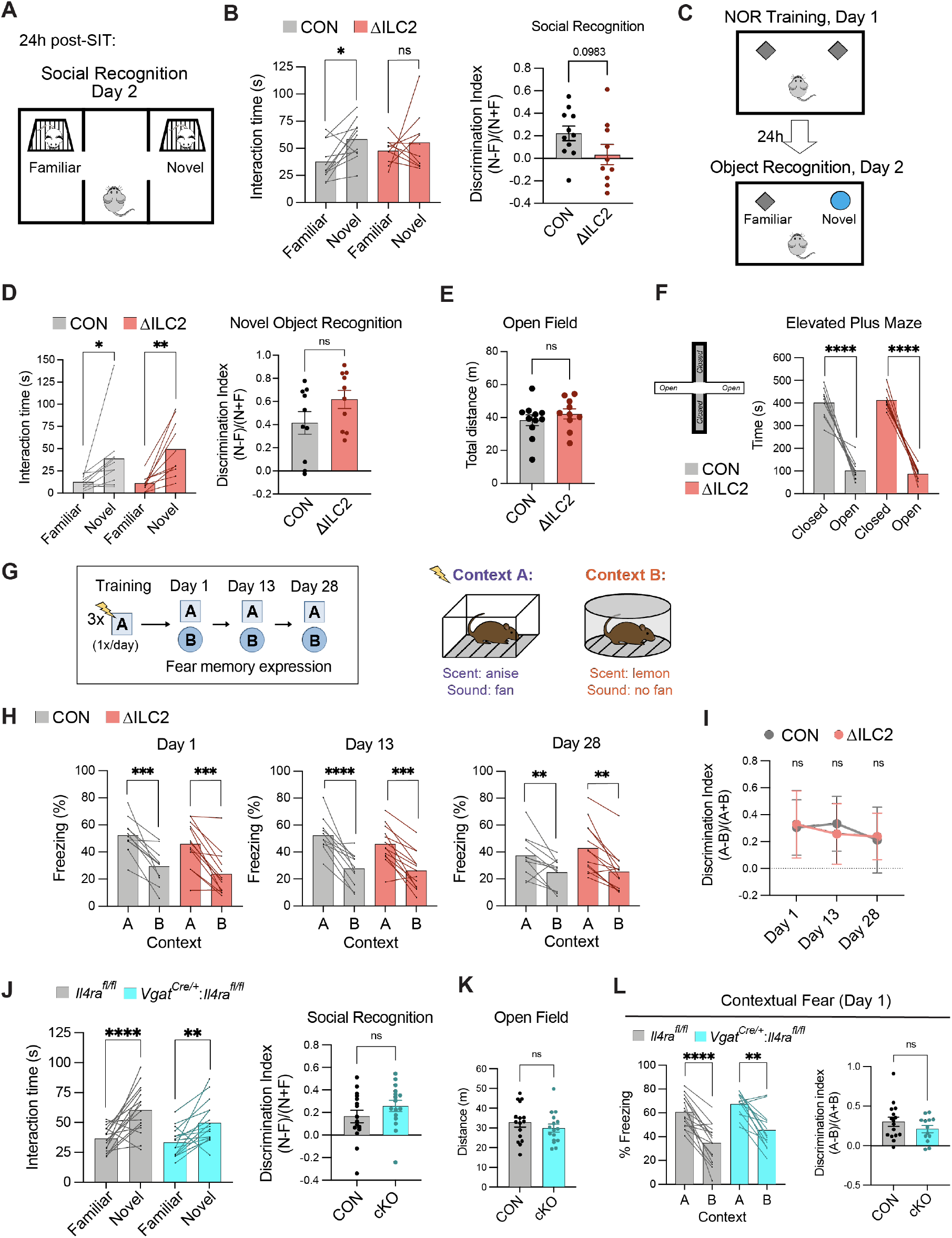
Additional behavioral assessments after ILC2 depletion or *Il4ra* conditional deletion from interneurons. **A)** Schematic of Crawley’s three-chamber assay of social novelty recognition (Day 2), 24 hours after assay of social interaction (day 1, shown in main Fig. 4A). **B) Left**: Interaction time spent with the Familiar vs. Novel mouse on Day 2 of the three-chamber social recognition assay for control vs. ΔILC2 mice. (Two-way ANOVA, Šídák’s multiple comparisons test). **Right**: The social recognition Discrimination Index for control vs. ΔILC2 mice calculated as the difference in interaction time with Novel (N) and Familiar (F) cup, over total interaction time ((N-F)/(N+F); Unpaired t test). **C)** Schematic of the novel object recognition assay (NOR). **D) Left**: Interaction time spent with the novel vs. familiar object for control vs. ΔILC2 mice (Two-way ANOVA, Šídák’s multiple comparisons test). **Right**: The novel object Discrimination Index for control vs. ΔILC2 mice calculated as the difference in interaction time with novel (N) vs. Familiar (F) object, over total interaction time ((N-F)/(N+F); Unpaired t test). **E)** Total distance traveled by controls and ΔILC2 mice in the open field assay (Unpaired t test). **F) Left**: Schematic of the elevated plus maze apparatus. **Right**: Time spent in closed arms vs. open arms by controls and ΔILC2 mice (Two-way ANOVA, Šídák’s multiple comparisons test). **G)** Schematic of the contextual fear discrimination assay. Mice were conditioned to a context with 1 footshock per day (context A) for 3 days, and then exposed to the conditioned context followed by an unconditioned but partially similar context (context B). Freezing was quantified in each context at 1, 13, and 28 days post-training. **H)** Percent of time spent freezing in the conditioned fear context A vs. unconditioned context B during retrieval tests at day 1, 13, and 28 post-training. **I)** The Discrimination Index calculated as the difference in freezing time in Context A and Context B, over total freezing time, ((A-B)/(A+B)) for controls and ΔILC2 mice on retrieval days. Dots represent group mean ± SD (Twoway repeated measures ANOVA, Šídák’s multiple comparisons test). **J) Left:** Interaction time spent with the familiar vs. novel mouse on Day 2 of the three chamber social recognition assay by control vs. interneuron *Il4ra* cKO mice (Two-way ANOVA, Šídák’s multiple comparisons test). **Right:** The Discrimination Index for social recognition calculated as the difference in interaction time with novel (N) vs. Familiar (F) cup, over total interaction time, for control vs. cKO mice ((N-F)/(N+F); Unpaired t test). **K)** Total distance traveled in the open field assay by controls and interneuron *Il4ra* cKO mice (Unpaired t test). **L)** Mice were conditioned to a context with 1 footshock per day (context A) for 3 days, and then exposed to the conditioned context followed by an unconditioned but partially similar context (context B). **Left**: Freezing was quantified in each context at 1 day post-training (Two-way ANOVA, Šídák’s multiple comparisons test). **Right**: The Discrimination Index was calculated as the difference in freezing time in Context A and Context B, over total freezing time on the retrieval day ((A-B)/(A+B); n=16 vs. 13 mice/genotype; Unpaired t test). For ILC2-deficient experiments: n = 11 vs. 10 mice/genotype. For interneuron *Il4ra* cKO experiments: n=16 mice/genotype, except where indicated above. Dots represent mice, except in panel I. Data are means with lines connecting paired data for each mouse (except I) and mean ± SEM. Statistics: *p < 0.05, **p < 0.01, ***p < 0.001, ****p < 0.0001.

## Acknowledgements

We are grateful to the A.V. and A.B. Molofsky labs, R.M. Locksley, and J.L. Rubenstein for helpful comments on the manuscript. We thank R.M. Locksley for generous sharing of mouse strains including *Il5^Cre-RFP^, R26R^DTA^, Il4ra^flox^* and *Il4ra^KO^;* M. Khierbeck, V. Sohal, and J. Crawley (UC Davis) for intellectual guidance and resources in behavioral analyses; K. Bender for electrophysiology guidance. We thank the UCSF Laboratory for Cell Analysis Core, the UCSF Parnassus Flow Core (RRID:SCR_018206 supported in part by Grant NIH P30 DK063720 and by the NIH S10 Instrumentation Grant S10 1S10OD021822-01), UCSF Biological Imaging and Development Core (BIDC), and UCSF Laboratory Animal Resource Center for instruments and services.

## Authors contributions

Conceptualization, J.J.B., A.V.M, and A.B.M.; Methodology, J.J.B., A.V.M, and A.B.M. Investigation, J.J.B., S.E.T., M.W.D. J.O.C., N.M.M., L.C.D., I.D.V. and C.C.E.; Writing – Original Draft, J.J.B., A.B.M. and A.V.M.; Writing – Review & Editing, all co-authors; Funding Acquisition, A.V.M, and A.B.M.; Resources, A.V.M, and A.B.M.; Supervision, A .V.M, and A.B.M.

## Data and materials availability

Supplement contains additional data. All data needed to evaluate the conclusions in the paper are present in the paper or the Supplementary Materials. RNA sequencing data is available through GEO #[submission in progress].

## Supplementary Materials

**Supplemental Table 1: Cluster defining genes for all meningeal cells scRNASeq:** Differential gene expression analysis for P14 meninges single cell RNA-seq clustering, related to Fig. 1B and S1D.

**Supplemental Table 2: Cluster defining genes for meningeal lymphocyte scRNASeq:** Differential gene expression analysis for non-B lymphocytes (cluster 9 in Fig. 1B) from P14 meninges single cell RNA-seq clustering, related to Fig. 1C and 1D.

## EXPERIMENTAL MODELS AND SUBJECT DETAILS

### Mice

All mouse strains were maintained in the University of California, San Francisco specific pathogen–free animal facility, and all animal protocols were approved by and in accordance with the guidelines established by the Institutional Animal Care and Use Committee and Laboratory Animal Resource Center. Littermate controls were used for all experiments when feasible, and all mice were backcrossed > 10 generations on a C57BL/6 background unless otherwise indicated. The mouse strains used are described in the Key Resources Table and as referenced in the text. All experiments, including behavioral analyses, incorporated animals of both sexes in approximately equal numbers. All experiments of n > 5 were analyzed for sex-specific trends, and none were evident unless specifically noted. For experiments using inducible conditional alleles, tamoxifen (Sigma, T5648) was dissolved at 10 mg/ml in corn oil (Sigma-Aldrich, C8267) shaking at 37°C overnight. For conditional deletion of IL-4Rα in myeloid cells (*Cx3cr1^CreER/+^*), mice were intragastrically injected with tamoxifen (50 μg/g body weight) at P1,3,5. For conditional deletion of IL-4Rα in microglia (*P2ry12^CreERT^*), mice were intragastrically/intraperitoneally injected with tamoxifen (50 μg/g body weight) at P3,5,7 and 100 μg at P11.

## METHODS DETAILS

### Meningeal dissociation

To prepare single cell suspensions for flow cytometric analysis and sorting, mice were deeply anesthetized and perfused transcardially with cold PBS. Horizontal cuts were made to remove the skullcap which was placed in ice cold FACS Wash Buffer (FWB; 0.1M DPBS (pH 7.4)/0.05% NaN3 /3% FBS). Under a dissecting microscope, the dural meninges were quickly removed by gentle scraping using fine forceps and pulled away in an intact sheet. Meninges were kept in ice cold cRPMI (RPMI 1640 supplemented with 10% (vol/vol) FBS, 1% (vol/vol) HEPES, 1% (vol/vol) Sodium Pyruvate, 1% (vol/vol) penicillin-streptomycin) until all samples were isolated. To obtain single cell suspensions, meninges were transferred to cRPMI containing 40μg/mL DNAse I and 0.2 wU/mL Liberase TM in 1 mL of cRPMI and incubated at 37°C with shaking at 220 RPM for 30 min. After gentle trituration, meninges and solution were passed through a 70-μm cell strainer; remaining tissue was firmly pressed through the filter in a circular motion using a wide tip plastic plunger from a 1 ml syringe. The strainer was flushed with additional FWB and collected into a 15 ml falcon tube. Cells were pelleted at 300 RCF for 7 min at 4°C and resuspended in FWB for downstream applications.

### Flow cytometry sample preparation

For cell sorting for single cell RNA sequencing, meningeal cell suspensions were incubated with Fc block (2.4G2) and stained with the following antibody solution in FWB (45 min at 4°C): CD45-BUV395, CD31-AF488, CD11b-PB, PDGFRa-APC, gp38-PECy7, DAPI (Antibody details in Key Resources Table). Cells were washed in FWB and fluorescence activated cell sorting was performed on a BD FACS Aria Fusion. Cells were gated for viable single cells and were enriched for lymphocyte and stromal populations by sorting CD45^+^CD11 b- (non-myeloid immune cells) and gp38^+^CD31- (stromal cells) separately, and adding each of those populations to 25% of total cells (1:1:2 ratio of lymphocytes: stroma: total viable populations, **Fig. S1A).** Genotypes of mice sequenced were IL-33 deficient (*Il33^mcherry/mcherry^*) or heterozygous controls (*Il33^mcherry/+^*). Meninges from both sexes were pooled by genotype to obtain technical replicates and sufficient numbers of cells for sorting. *Il33^wt/mCherry^* controls included 5 females and 7 males, and *Il33^mCherry/mCherry^* knockouts included 7 females and 8 males.

For immunophenotyping over development, flow cytometric analyses were performed. For cytokine reporter tracking, cell suspensions were incubated with a viability dye (Zombie-NIR, 20 minutes), followed by Fc block (2.4G2) and stained with the following antibody solution in FWB (45 min at 4°C): CD45-BUV395, CD3e-AF700, CD4-BV711, CD8a-BV785, Thy1.2-BV421, CD11b-BV605, NK1.1-BV650, CD19-PEDazzle 594, CD11c-PeCy7, CD25-PerCPCy5.5, Hu-CD4-APC (Antibody details in Key Resources Table), followed by wash and resuspension in FWB prior to analysis. Intracellular stains were combined with intravenous labelling of blood circulating cells. Three minutes prior to euthanasia, mice were administered intravenous CD45.2-APCCy7 to label blood circulating cells. Cell suspensions were incubated with a viability dye (Live/Dead Aqua, 20 minutes), followed by Fc block (2.4G2) and surface marker staining with the following antibody solution in FWB (45 min at 4°C): CD45-BUV395, CD3e-AF700, CD4-BV711, CD8a-BV785, Thy1-BV421/PB, CD11b-BV605, TCR γ/δ- PerCPCy5.5, NK1.1-BV650, CD19-PEDazzle 594 (Antibody details in Key Resources Table). Cells were washed in FWB and fixed and permeabilized (60 min at RT) using the FoxP3/transcription factor staining buffer set (eBioscience), followed by intracellular staining with the following antibody solution in 1x PermBuffer (eBioscience) supplemented with 5% rat serum): Rorγt-APC, EOMES-FITC, Tbet-PeCy7, GATA3-PE (60 min at 4°C). Cells were washed two times in 1x PermBuffer and resuspended in FWB prior to analysis. Flow cytometric analysis was performed on a BD Fortessa and data analysis was performed using FlowJo™ software (BD).

### Single cell RNA sequencing and analysis

Approximately 35,000 sorted cells of each sample were loaded per well of the Chromium Chip B (10X Genomics) and libraries were generated using the Single Cell 3’ reagent kit v3 (10X Genomics) by the UCSF Institute for Human Genetics Core, according to the manufacturer’s instructions. Sequencing was performed on the NovaSeq 6000 Illumina sequencing platform reaching a mean reads per cell of 48,729 (*Il33^mcherry/+^*) and 51,733 (*Il33mcherry/mcherry*).

Sequenced samples were processed using the Cell Ranger 3.0.2 pipeline, aligned to the GRCm38 (mm10) mouse reference genome, and further analyzed using R (v 3.6.1) and Seurat (v 3.1.5, Satija et al., 2015). Each data set was individually processed filtering cells within the following thresholds: mitochondrial content <15%, number of genes >500 and <7500 and hemoglobin content <0.1%. Following quality control filtering, 5950 cells and 6594 cells, respectively, were used for analysis and integration. Samples were individually SCTransform normalized (regressed to mitochondrial content and cell cycle score (G1 and S-phase genes: (G1) *Tacc3, Hmgb3, Hmmr, Nde1, Gtse1, Spag5, Kif22, Tpx2, Birc5, Cdc25b, Ube2s, Cdca3, Adgrg6, Ttk, Brd8, Sfpq, Cdc20, Nek2, Cenpf, Gpsm2, Cks2, Top2a, Ccnb1, Cdca8, Troap, Espl1, Ckap2, Nusap1, Kif23, Kif11, Cenpe, Kif2c, Lbr, Ccna2, Mki67, Bub3, Ccnb2, Cdc25c, Ccnf, Hmgb2, Pttg1, Plk1, Bub1, Ube2c, Kpna2, Gm10184, Arhgap11a;* (S) *Mcm2, Mcm6, Ung, Cdc6, E2f1, Ccne1, Pask, Pcna, Dtl, Hspb8, Rfc4, Slbp, Chaf1a, Exo1, Ccne2*) and integrated based on 3000 anchoring features using the SCTransform workflow (https://satijalab.org/seurat/v3.2/integration.html). Following integration, 30 principal components (PCs) were calculated and the first 20 PCs were used for clustering at a resolution of 0.5. The clusters were identified based on expression of canonical markers. Non-B lymphocytes (cluster9) were further subclustered from the integrated dataset resulting in a total of 477 cells (224, and 253 cells from respective sample) and re-clustered using 2000 variable features and 20 PCs at a resolution of 0.3. Cells were identified as ‘‘female’’ or ‘‘male’’ based on their expression of the gene *Xist;* any cells expressing one or more counts of Xist were labeled female, while all others were labeled male. We then used these assignments to bioinformatically distinguish replicates for a total of two replicates for genotype. Differential expression for each cluster shown in plots Fig. 1B and 1C were calculated using the FindAllMarkers function in Seurat on genes expressed in at least 25% of the cells in that cluster. P-values were calculated on upregulated genes using a Poisson test on genes with >0.25 natural log (base e) fold change. The heatmap was created in Seurat using the top 10 markers per cluster based on log fold change.

### Whole mount meningeal preparation and immunostaining

For whole mount meninges imaging, neonatal (P1 or P14) mice were perfused transcardially with 5-10 mL of ice-cold 1X PBS followed by 5-10 mL of 4% (weight/volume) paraformaldehyde (PFA). Skullcaps containing dural/arachnoid meninges were separated from the brain/pial membrane and post-fixed in 4% PFA overnight at 4°C with shaking (90 rpm). Skullcaps/meninges were washed with 1X PBS and then incubated in Blocking/Permeabilization Buffer (0.3% Triton X-100, 5% fetal bovine serum (FBS), 0.5% bovine serum albumin (BSA), and 0.05% NaN3 in PBS) for at least one overnight at 4°C (90 rpm). Skullcaps/meninges were then incubated with primary antibodies in Staining Solution (0.15% Triton X-100, 7.5% FBS, 0.75% BSA, and 0.075% NaN3 in PBS) for three days at 4°C (90 rpm). Skullcaps/meninges were washed three times in PBS-T (0.15% Triton X-100) for 30 min each at 4°C (90 rpm) and incubated with secondary antibodies in Staining Solution overnight at 4°C. Skullcaps/meninges were washed three times in PBS-T for 30 min each and then incubated with directly conjugated antibodies in Staining Solution overnight at 4°C (90 rpm). Skullcaps/meninges were again washed three times in PBS-T for 30 min each and then stained in DAPI (2 μg/mL in PBS) for 30-60 min at 4°C (90 rpm), and rinsed in 1X PBS. Meninges were then dissected out of skullcaps under a dissecting microscope in a PBS bath and whole mounted onto glass slides. Residual PBS was wicked away and replaced with RIMS (Refractive Index Matching Solution: 80% (w/v) Histodenz in 1X PBS, 0.01% NaN3, 0.1% Tween20). Meninges were covered with a glass coverslip, sealed, and allowed to clear at least one day prior to imaging.

For meningeal imaging, the following antibodies were used: rat anti-CD31 (clone MEC13.3), goat antirat AF647, and mouse anti-smooth muscle actin (αSMA)–AF488 direct conjugate (clone 1A4). See Key Resources Table for antibody details and dilutions.

### Brain immunostaining

For all immunostaining experiments, mice were perfused transcardially with 10 mL of ice-cold 1X PBS followed by 10 mL of 4% (weight/volume) paraformaldehyde. Brains were post-fixed in 4% PFA at 4°C for 4-5 hours unless otherwise noted, and cryoprotected in 20% sucrose solution for a minimum of 2 days. Brains were flash frozen and sliced in 40 um-thick coronal sections on a HM440E freezing microtome (GMI Instruments). In some cases, brains were embedded in OCT and stored at −80°C until sectioning in 40 um-thick coronal sections on a CryoStar NX70 Cryostat (Thermo Fisher). For RNAscope experiments, brains were fixed in 4% PFA overnight, cryopreserved and embedded, sliced in 14 um-thin sections and slide-mounted.

Free floating 40 um-thick sections were washed three times with PBS prior to antigen retrieval or blocking to remove cryopreservative. For synaptic stains, antigen retrieval was performed in 0.01M sodium citrate pH 6 (95°C for 10 min), cooled at room temperature for 5 min in solution, and washed three times in PBS. Sections were blocked for one hour at room temperature in Staining Solution (0.4% TritonX-100, 5% normal goat serum, 1X PBS), then stained overnight with primary antibody mix in Staining Solution (rocking at 4°C). Sections were washed three times in PBS-T (0.05% Triton X-100 in 1X PBS) and stained with secondary antibody mix in Staining Solution for 1.5 hours (rocking at room temperature). Sections were washed three times in PBS-T, counterstained with Hoechst or Dapi, and mounted onto slides with Fluoromount-G (SouthernBiotech #0100-01).

For optimal synapse staining, the following measures were always taken: Antigen retrieval was performed; Staining Solution was made from high purity 10% Triton X-100 Surfact-Amps detergent (ThermoFisher #28314); Antibody mixes were centrifuged at 15,000 RCF for 3 min in 1.5 ml tubes to remove any aggregates. The following primary antibodies were used: mouse anti-Gephyrin, rabbit anti-VGAT, rabbit anti-PSD95, guinea pig anti-VGLUT2, chicken anti-NeuN, rabbit anti-IBA1, rabbit anti-GAD67, and mouse anti-Parvalbumin. The following secondary antibodies were used: Goat antimouse AF647, Goat anti-rabbit AF555, Goat anti-guinea pig AF647, and Goat anti-chicken AF488. See Key Resources Table for antibody details and dilutions.

### RNAscope *in situ* hybridization

FISH experiments were performed using the RNAscope Multiplex Fluorescent Reagent Kit v2 assay (ACD Bio) as described by the manufacturer for fixed-frozen tissue, except tissue was not baked prior to tissue dehydration, antigen retrieval was performed at 80°C, and protease III treatment was reduced to 20 min. Brains were embedded in OCT following 20% sucrose treatment, frozen at −80°C for a minimum of 1 day and then sliced in 14 um-thick coronal sections on a CryoStar NX70 Cryostat (Thermo Fisher) before being mounted on slides for downstream RNAscope processing. Hybridization was performed with the *Mm-Il4ra-C1* probe (ACD Bio, #520171) and TSA Plus Cyanine 3 or Cyanine 5 dye (Perkin Elmer) to detect the receptor subunit transcripts. For immunohistochemical labeling with antibodies following the RNAscope assay, tissues were incubated with blocking and antibody solutions as described above immediately after developing the HRP-TSA Plus signal and washing three times.

### Confocal imaging and Image analysis

#### Quantification of meningeal ILC2s

Whole mounted meningeal tissues were imaged with a Nikon A1R laser scanning confocal microscope (405, 488, 561, and 650 nm laser lines) using a 16X/0.8 NA Plan Apo water-immersion objective (512 resolution, 1 frame/s, Z step-size = 4um). Z-stack images were rendered into three dimensions and quantitatively analyzed using Bitplane Imaris v9.5.1 software (Andor Technology PLC, Belfast, N. Ireland). 3D reconstructions of ILC2s were generated and quantified using the Imaris surface function on *Il5*-tdTomato^+^ cells, thresholded on signal intensity, volume, and sphericity, as described previously (Dahlgren et al., 2019; Cautivo et al., 2022).

#### Quantification of synapses

For synaptic puncta quantification, brain sections were imaged on a Zeiss LSM 800 confocal microscope with a 63X/1.4 NA Plan-Apochromat oil-immersion objective. Image acquisition settings: Field size: 101.4 μm x 101.4 μm (10,281.96 μm); Pixel size: ~0.08 μm (1200×1200); Scan speed 4; Averaging: 4; 16 bit; Power, master gain, and digital gain were adjusted for individual experiments and kept consistent within an experiment. Single optical sections were acquired at a consistent depth of 5 μm below the surface to reduce variability of signal. We conducted our analysis in layer 4 of S1 somatosensory cortex medial to the barrel field (**Fig. 3D**) in order to achieve a highly consistent subregion, to avoid anatomical variability (e.g. due to the barrel septa), and due to enrichment of thalamocortical synapses for VGLUT2 analysis. Each image contained only L4 as determined by DAPI nuclei density and NeuN cell bodies if included. For IL-13 i.c.v injection experiments, the same region was quantified at ~100-250 μm from the injection site. Technical replicates consisted of 2 images per brain section, 3 sections per mouse (total 6 images); at least 4 biological replicates per genotype or condition were collected, and littermates were used within independent experiments, unless otherwise noted.

The ImageJ2 plugin PunctaAnalyzer was used to quantify the numbers of synaptic puncta and the colocalization of presynaptic and postsynaptic marker channels, as described previously (Stogsdill et al., 2017; Vainchtein et al., 2018). Pre- and postsynaptic marker pairs for inhibitory (VGAT and Gephyrin) and excitatory thalamocortical (VgluT2 and PSD95) synapses were used. Background subtraction (rolling ball size 50 pixels) was applied, and threshold set to the max tail of the intensity histogram per channel for each image. Puncta counts were normalized to the mean of controls within each experiment.

#### Quantification of perisomatic and axon initial segment synapses

Z stacks were acquired with a Zeiss LSM800 confocal microscope using a 40X/1.0 NA Plan-Apochromat oil objective. For perisomatic synapse analysis, single optical sections containing multiple NEUN^+^ soma were analyzed in FIJI. Neurons with cross sections at the center of the soma were quantified. An ROI was manually drawn around a NEUN^+^ soma to measure perimeter and VGAT^+^ puncta contacting this perimeter were counted. Ten neurons from 2-3 images per mouse were quantified. For AIS synapse analysis, surface reconstructions were rendered from Z stacks using Imaris software (v9.8) Cell:Vesicle model. VGAT^+^ spots in contact with the ANKG^+^ surface were quantified. Only complete AIS objects in the volume were selected for analysis of length and puncta contacts. Approximately 25-28 AIS from two Z stacks per mouse were quantified.

#### Quantification of inhibitory neurons

Brain sections were stained for GAD67, Parvalbumin (PV), and NeuN. Sections were imaged on a Zeiss Z2 fluorescent microscope with Apotome using a 20X/0.8 NA Plan-Apochromat air objective. A tiled field was imaged containing S1 somatosensory cortex L1-L6. Images were analyzed in FIJI by manual drawing of ROI containing each cortical layer and counting GAD67^+^ or PV^+^ soma. Density was calculated by dividing number of cells counted over the area of the ROI. All soma were confirmed to be NeuN^+^. Two images from two brain sections were averaged per animal, from 2 experiments; littermates were used within independent experiments.

#### Quantification of RNAscope FISH

Hybridized and immunostained slides were imaged on a Zeiss LSM 800 confocal microscope with a 20X/0.8 NA Plan-Apochromat air objective. A tiled Z stack field was imaged containing S1 somatosensory cortex (4-5 μm thick, Z-step size 0.5 μm). Maximum intensity Z projections were created in FIJI. For layer puncta density quantification, an ROI was drawn for each cortical layer and particle analysis was performed to obtain the number of puncta per layer ROI. For cell type quantification, total numbers of microglia (IBA1^+^ soma), neurons (NeuN^+^ soma), and interneurons (*Vgat^Cre^*;Ai14^+^ soma) were manually counted within an ROI. Number of microglial and neuronal soma positive for *Il4ra* transcripts (at least 2 puncta localized to the nucleus) were counted. Density was calculated as counts per area of the ROI analyzed. One tiled image per section, 2 sections per mouse, for two mice were counted per condition.

### Slice Preparation and Patch-Clamp Electrophysiology

Mice were deeply anesthetized with isoflurane and euthanized. The brain was removed and 250 μm-thick coronal slices were prepared in ice-cold sucrose cutting solution (234 mM sucrose, 2.5 mM KCl, 1.25 mM NaH2PO4, 10 mM MgSO4, 0.5 mM CaCl2, 26 mM NaHCO3, and 10 mM glucose, equilibrated with 95% O2 and 5% CO2, pH 7.4) using a Leica VT1200 microtome (Leica Microsystems). Sections containing Layer IV barrel cortex were incubated for an hour at 32-34°C and then at room temperature in artificial cerebrospinal fluid (aCSF; 126 mM NaCl, 2.5 mM KCl, 1.25 mM NaH2PO4, 1 mM MgCl2, 2 mM CaCl2, 26 mM NaHCO3, and 10 mM glucose, equilibrated with 95% O2 and 5% CO2, pH 7.4). Recording electrodes were made from borosilicate glass with a resistance 3–5 MU when filled with cesium chloride intracellular solution (129 mM CsCl, 10 mM HEPES, 10 mM EGTA, 5 mM QX-314 Cl, 4 mM MgATP, 2 mM MgCl2, pH adjusted to 7.35 with CsOH; 286 mOsm). Series resistance was monitored in all recordings, and the recordings were excluded from analysis if the series resistance was > 25 MOhm or varied by more than 25%. Recordings were obtained using a MultiClamp 700B microelectrode amplifier (Molecular Devices, Sunnyvale, CA), digitized using Digidata 1550B (Molecular Devices), and acquired at 20 kHz using pClamp 10 software (Molecular Devices). Recordings were performed in voltage-clamp mode at a holding potential of −65 mV and obtained from visually identified spiny stellate cells within the barrels of the barrel cortex. In the presence of aCSF supplemented with 0.5 μM tetrodotoxin (TTX), 20 μM DNQX, and 50 μM APV, miniature inhibitory post-synaptic currents (mIPSCs) were isolated and recorded for 10 minutes. Recordings were analyzed using ClampFit (Molecular Devices) and Wdetecta software (https://huguenard-lab.stanford.edu/wdetecta.php). Experimenter was blinded to genotype.

### Behavioral Assays

Experiments were conducted in 8-14 week old mice, experimenter was blinded to genotype throughout data collection and analysis. ILC2-deficient and control mice underwent a series of assays in order in two independent cohorts: Open Field Test, Novel Object Recognition, Object Place Recognition, Social Interaction Test, Social Recognition Memory test, Elevated Plus Maze, and Contextual Fear Conditioning. Inhibitory neuronal conditional knockout and control mice underwent the same assays in order except Elevated Plus Maze. To acclimate mice to the experimenter prior to the behavioral testing, mice were gently handled in their home cage for 8 min per day for 3 days. Testing occurred over 13 days. Mice were habituated to the testing room 1-2h prior to each assay. ANY-maze software (Stoelting Co.) was used in all assays, except where noted, to video record and track mouse movement using automated object detection and zones customized to each apparatus. In one instance following unblinding, data from two male ILC2-deficient mice and one control was excluded from analysis due to discovery of inadvertent cohousing with a female mouse for three weeks prior to testing. Tracking plots were checked for accuracy in all assays and retracked when necessary. Manual scoring was performed for a subset of mice per assay to validate accuracy.

#### Social interaction test

Sociability and preference for social novelty were measured using Crawley’s Three-Chamber test in adult mice as described previously (Nadler et al., 2004; Silverman et al., 2010).

One day before the test, mice were habituated for 12 min to the three-chamber open field with equally sized and connected chambers. To test sociability, on test day 1, an ovariectomized female mouse (8-12 weeks) was placed in a wire cup in the left or right chamber (Social cup) and an empty wire cup was placed in the opposite chamber (Empty cup). The experimental mice were placed in the center chamber and allowed to freely explore the three-chamber open field for 10 minutes. Their exploration was recorded and tracked using ANY-maze software. The location of the social cup was alternated in the left or right chamber between mice. The chamber was cleaned thoroughly after each mouse. “Interaction time” was counted when the nose of the test mouse entered the zone drawn immediately (~1 cm) around the empty or social cup.

#### Social recognition memory test

Twenty-four hours after the Social Interaction Test, mice were tested for social memory. The previously presented ovariectomized female from day 1 was placed in the same wire cup (Familiar cup) in the same chamber, and a novel ovariectomized female was placed in the previously empty opposite cup (Novel cup). The experimental mice were placed in the center chamber and allowed to freely explore the three-chamber open field for 10 minutes. Their exploration was recorded and tracked using AnyMaze software.

#### Open field test

Mice were introduced to an open field chamber and were assessed over a 10 minute exposure. Their exploration was recorded and tracked using AnyMaze software to obtain distance traveled, time spent freezing, and time in the center zone of the field.

#### Novel object recognition

Novel object recognition was performed as previously described (Antunes and Biala, 2012). The Open Field Test (OFT) served as habituation for novel object recognition test. The day after the OFT, mice were placed in the same open field apparatus that now contained two identical objects and allowed to explore for 7 min. Twenty-four hours later, mice were tested for object memory and presented with one object from the prior day (Familiar) and one new object (Novel). Location of the novel object was alternated between mice. Exploration was recorded and tracked using AnyMaze software. “Interaction time” was counted when the nose of the test mouse entered the zone drawn immediately around the objects. Objects used in the study were small glass jars of different shape and size.

#### Elevated plus maze

Mice were introduced to an elevated plus maze and were assessed over a 10 minute exposure. Open arms of the maze were identified as the two opposing arms without walls, and closed arms of the maze were identified as the two opposing arms with walls. Their exploration was recorded and tracked using AnyMaze software to obtain time spent in the open and closed arms of the maze.

#### Contextual fear conditioning

Conditioned fear was elicited by administering a mild footshock (0.75 mA) following a 3 minute exposure to an array of contextual cues (conditioning chamber, chamber lights, white noise, scent of anise extract) for 3 days. Retrieval of the fear memory was assessed by a 5 minute re-exposure of the animal to the conditioning context in the absence of shock, and freezing (the cessation of all movement outside of respiration) was interpreted as expression of fear memory. Video recordings were acquired and scored automatically in FreezeFrame (Actimetrics). Mice were habituated to transport and holding in a separate room for 1-2 hours prior to all conditioning or retrieval sessions and subjected to three days of handling prior to all behavioral tests. For assessment of fear discrimination, freezing was measured in a context similar to the conditioning context 2 hours later, but with the following variations: the chamber fan was turned off, the scent of lemon extract instead of anise extract was used, and a plastic divider was inserted to make the chamber walls circular and opaque. Freezing in the similar context was tested 2 hours following retrieval testing in the original conditioning context, and animals were rested in a holding room between sessions.

### ILC2 transfers

*In vivo* expansion and activation of ILC2s was performed in *Il5^Cre-RFP/+^;R26^TdT/+^* mice with 3 injections of 500 ng recombinant carrier-free murine IL-33 (BioLegend) per mouse as described previously (Cautivo et al., 2022); a final injection was given to activate cells 3-4 days prior to transfer. To obtain sufficient cell numbers, ILC2s were isolated from the lung. Donor mice were euthanized with CO2, transcardially perfused with 10 mL of ice cold 1X PBS, and lungs were collected into 1X Hank’s Balanced Salt Solution (HBSS). Lungs were cut into small pieces with an automated tissue dissociator (GentleMacs; Miltenyi Biotec) using the “lung1” program and then digested in 1X HBSS with 0.2 wU/mL Liberase TM (Roche) and 40 μg/mL DNase I (Roche) for 30 min at 37°C with shaking (200 rpm). Samples were homogenized with the GentleMacs using the ‘‘lung2’’ program, filtered through 70μm filters, washed, and subjected to red blood cell lysis (1X Pharm-Lyse solution; BD Biosciences). Single cell suspensions were stained and washed in FACS Wash Buffer (1X DPBS, 3% FCS, 0.05% NaN3) prior to sorting on a FACSAriaII. Lung ILC2s were sorted as viable, CD45^+^, lineage negative (CD8α- CD11b- CD11c- CD19- NK1.1- Ter119- SiglecF- CD3- CD4-) RFP^+^ (l5^Cre-RFP^;R26^TdT+^) cells and collected in complete RPMI (1X RPMI 1640 supplemented with 10% (v/v) fetal bovine serum, 1% (v/v) penicillin/streptomycin, 1% (v/v) Glutamax, 1 mM sodium pyruvate, 10 mM HEPES, 10 mM nonessential amino acids, and 55 μM β-mercaptoethanol). Cells were washed in normal saline and resuspended to 20,000 cells/μl before i.c.v. injection the same day. Saline or 20,000 ILC2s were transferred into ILC2-deficient (ΔILC2) recipient mice at P2,3, or 4 (bilateral i.c.v. injection, 0.5 μl per side). Recipient mice were euthanized at age P30 and brains were taken for histology.

### Stereotactic injections

All brain injections were performed with a Kopf stereotaxic apparatus (David Kopf, Tujunga, CA) and a microdispensing pump (World Precision Instruments) holding a Hamilton Syringe (model 701 RN, 10 μl) with a beveled glass needle (~50 mm outer diameter). For intraventricular injections into neonates (P2-4), mice were anesthetized by hypothermia on ice for 3 min. Bilateral i.c.v. injection of cell suspension or saline vehicle (0.5 μl per side) was performed (from lambda: 1.8 mm AP, +/-1 mm ML, −1.55 mm DV). Afterwards, pups were warmed on a heating pad until full recovery before returning to their home cage. For intraventricular injections into juveniles (P14), mice were anesthetized with 1.5% isoflurane at an oxygen flow rate of 1L/min, head-fixed with a stereotaxic frame (for juveniles, size P11), and treated with ophthalmic eye ointment. Fur was shaved and the incision site was sterilized with 70% ethanol and Betadine prior to surgical procedures. Subcutaneous 0.5% lidocaine was administered at the incision site and lack of reflex response was checked. Body temperature was maintained throughout surgery using a heating pad. A hole was drilled in the skull and 1.25 μl of recombinant carrier-free murine IL-13 (200 ng/μl, BioLegend) or saline was injected (from lambda: 3.7 mm AP, 1 mm ML, −1.75 mm DV) at a rate of 4 nL/sec. The needle was held in place for 5 minutes to allow diffusion and then slowly removed. The incision was closed and mice were allowed to fully recover with heat. Buprenorphine (Henry Schein Animal Health) was administered (0.1 mg/kg) according to approved protocols (briefly, mice were dosed prior to surgery, 4-8h later, and the next morning if needed by intraperitoneal injection).

### Bulk RNA sequencing of cortical microglia

Microglia were isolated from cortex, stained with CD45-FITC (1:100, BioLegend) and CD11b-APC (1:100, BioLegend), and sorted on an Aria III (BD) flow cytometer into RLT plus buffer (QIAGEN), as described previously (Galatro et al., 2017). RNA was isolated from 30000-60000 microglia per sample with the RNeasy^®^ Plus Micro kit (Qiagen). Quality and concentration were determined with the Agilent RNA 6000 Pico kit on a Bioanalyzer (Agilent). All samples had an RNA Integrity Number (RIN) >7. cDNA and libraries were made using the Ovation^®^ RNA-Seq System V2 kit (NuGen) and quality was assessed by Agilent High Sensitivity DNA kit on a Bioanalyzer (Agilent) and quantified by qPCR. Pooled libraries were RNA sequenced on an Illumina HiSeq 4000 paired-end for 125 cycles (PE125) yielding 50-70 million reads per sample.

### Bulk RNA sequencing analysis

Quality of reads was evaluated using FastQC (http://www.bioinformatics.babraham.ac.uk/projects/fastqc), all samples passed quality control, and reads were aligned to mm10 (GRCm38; retrieved from Ensembl, version November 2019) using STAR (version 2.5.4b) (Dobin et al., 2013) with ‘–outFilterMultimapNmax 1’ to only keep reads that map one time to the reference genome. Mapped reads were counted using HTSeq (version 0.9.0)(Anders et al., 2015) and DESeq2 package (version 1.24.0) (Love et al., 2014) was used to normalize the raw counts and perform differential gene expression analysis. GO analysis was conducted using the Metascape webpage (https://www.metascape.org) for genes from the comparison IL-13 versus PBS with adjusted p value < 0.01 and log fold change >1.

## QUANTIFICATION AND STATISTICAL ANALYSIS

Graphpad Prism 9.4.1 was used for most statistical analyses. Statistical tests are as described in text and figure legends. RNA-sequencing data was analyzed in R as described in the methods section above.

## REAGENTS AND RESOURCES

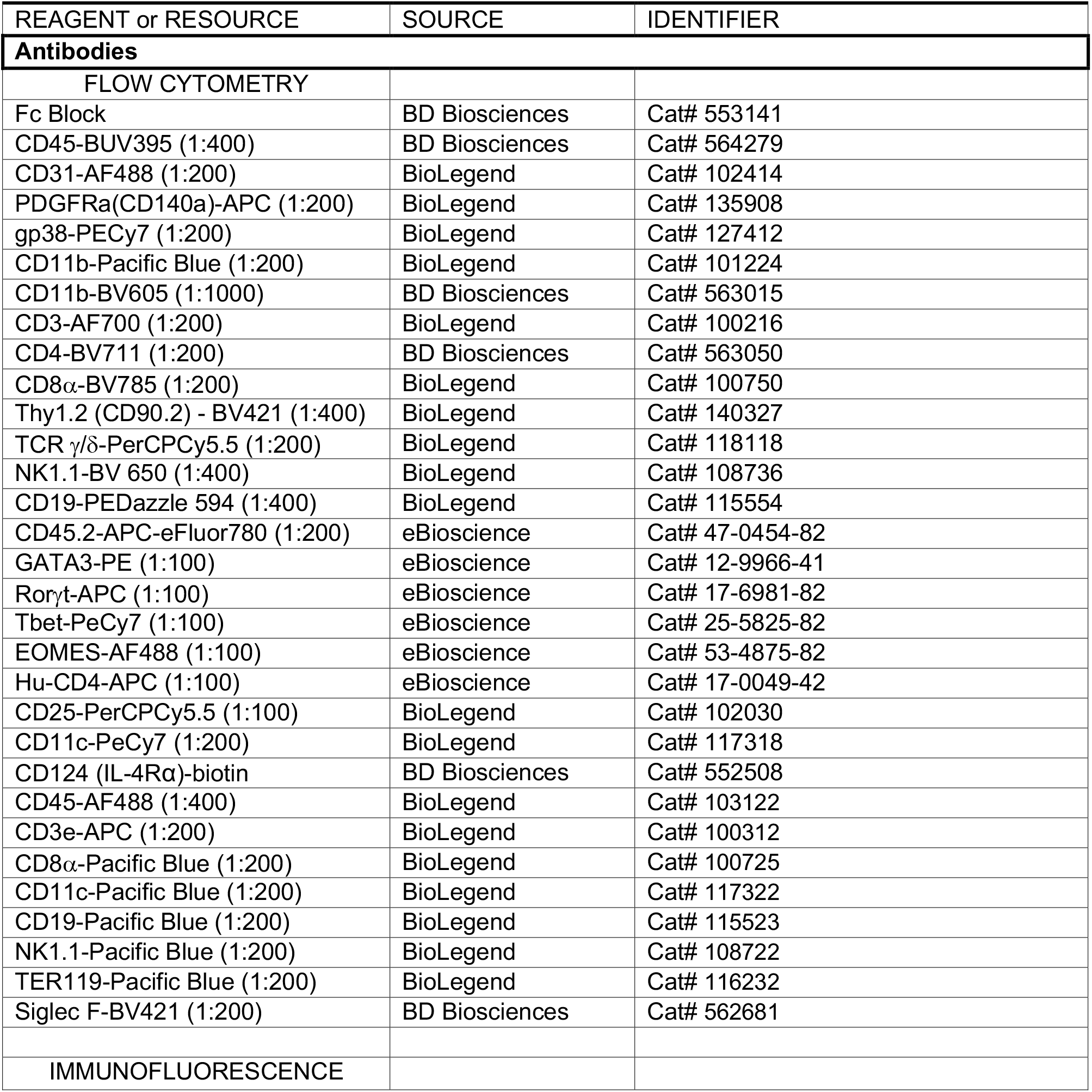
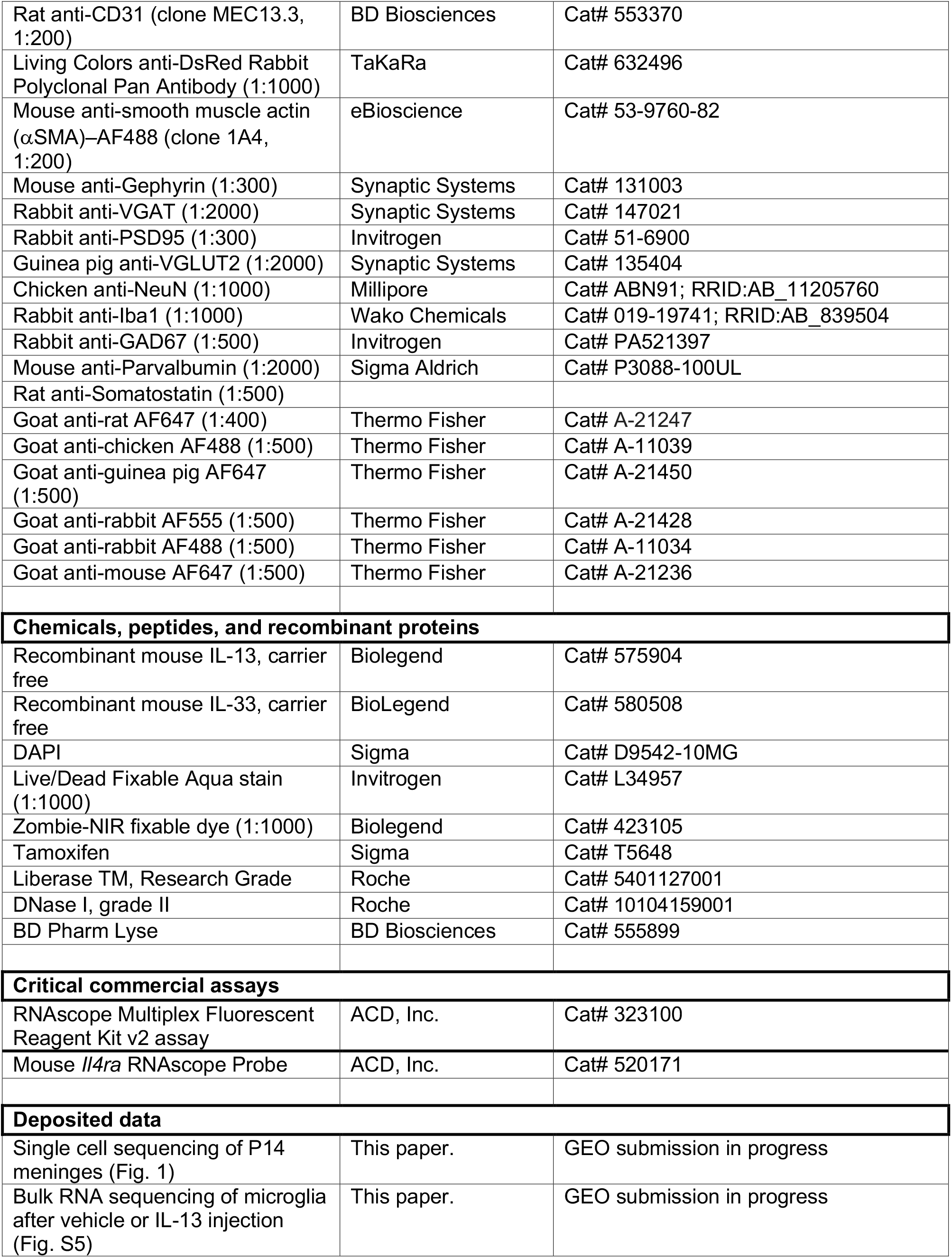
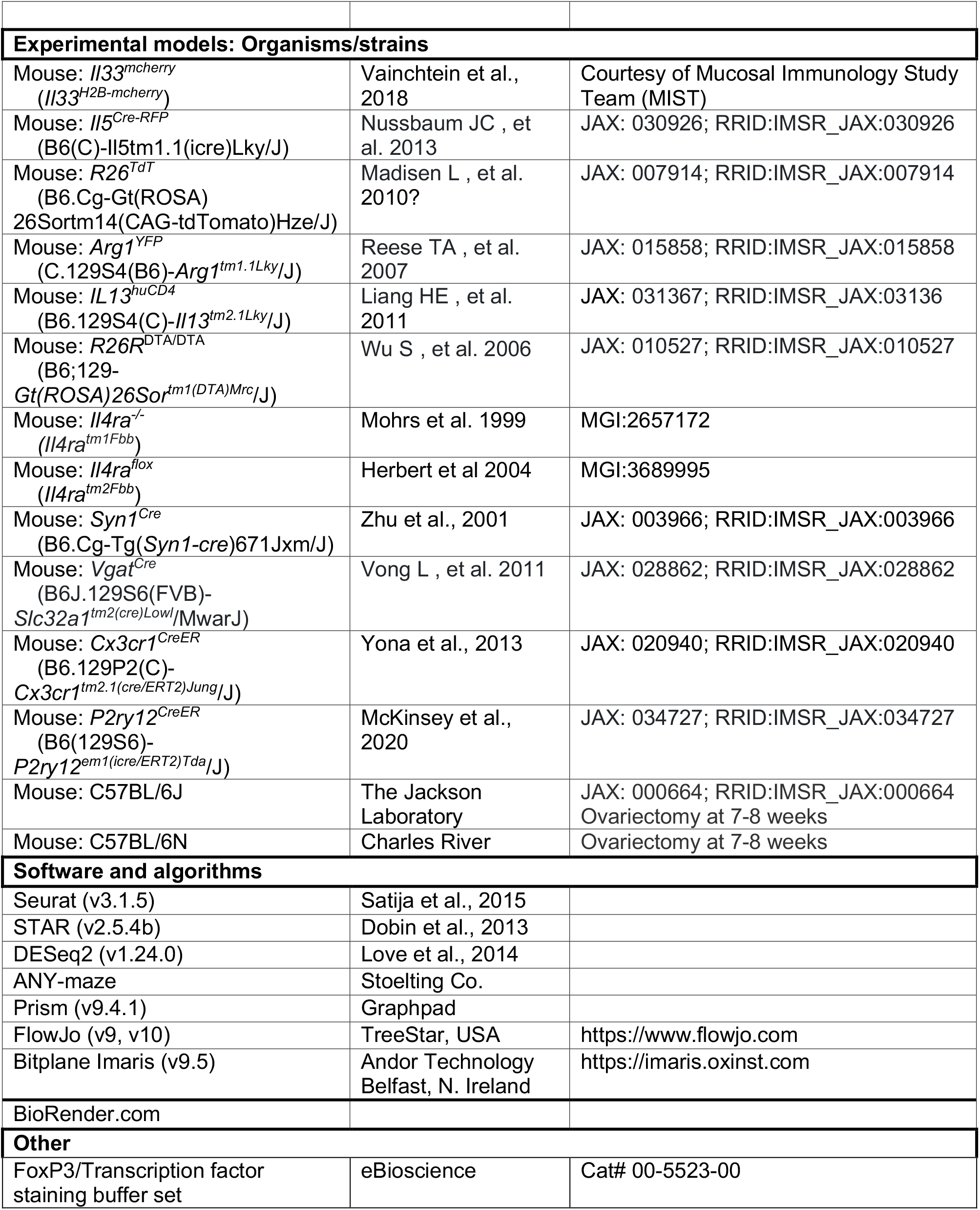

